# Reorganization of 3D Genome Structure May Contribute to Gene Regulatory Evolution in Primates

**DOI:** 10.1101/474841

**Authors:** Ittai E. Eres, Kaixuan Luo, Chiaowen Joyce Hsiao, Lauren E. Blake, Yoav Gilad

**Author notes:** Correspondence should be addressed to YG.

## Abstract

A growing body of evidence supports the notion that variation in gene regulation plays a crucial role in both speciation and adaptation. However, a comprehensive functional understanding of the mechanisms underlying regulatory evolution remains elusive. In primates, one of the crucial missing pieces of information towards a better understanding of regulatory evolution is a comparative annotation of interactions between distal regulatory elements and promoters. Chromatin conformation capture technologies have enabled genome-wide quantifications of such distal 3D interactions. However, relatively little comparative research in primates has been done using such technologies. To address this gap, we used Hi-C to characterize 3D chromatin interactions in induced pluripotent stem cells (iPSCs) from humans and chimpanzees. We also used RNA-sequencing to collect gene expression data from the same lines. We generally observed that lower-order, pairwise 3D genomic interactions are conserved in humans and chimpanzees, but higher order genomic structures, such as TADs, are not as conserved. Inter-species differences in 3D genomic interactions are often associated with gene expression differences between the species. To provide additional functional context to our observations, we considered previously published chromatin data from human iPSCs. We found that inter-species differences in 3D genomic interactions, which are also associated with gene expression differences between the species, are enriched for both active and repressive marks. Overall, our data demonstrates that, as expected, an understanding of 3D genome reorganization is key to explaining regulatory evolution.

## Introduction

Over the last several decades, a growing body of evidence indicated that variation in gene regulation plays a key role in phenotypic divergence between species [1–7]. Inferring the causal relationship between inter-species regulatory differences and phenotypic differences between species remains challenging, but compelling examples of regulatory adaptations have been published in a large number of species, including primates [8–13]. The molecular mechanisms that underlie regulatory adaptation have also been the focus of much research. In mice, flies, yeast, and primates, studies have revealed that expression divergence between species is often largely driven by mutations or epigenetic modifications within cis-regulatory elements (CREs), rather than *trans* elements (e.g. transcription factors) [5, 8-12, 14-15]. This makes intuitive sense, because whereas transcription factors may operate broadly across multiple functional contexts and the entire genome (affecting many genes), CREs often have a much more specific functional outcome [3, 16].

The ability to measure epigenetic marks, chromatin structure, and other functional genomic data has enabled us to identify and classify CREs into different types of regulators with distinct effects on gene expression (e.g. enhancers, silencers, insulators) [17–19]. Despite significant advances in our ability to identify and predict the functional role of CREs, we still lack a comprehensive characterization of the functional relationships between CREs and the genes they regulate. In other words, we still often do not know which genes are regulated by which CREs, or when and how often these relationships change. Connecting CREs to their target genes is crucial for understanding how regulatory architecture changes in response to different spatial, temporal and organismal contexts [15, 17-22]. Ultimately, the effects of CREs on gene expression are likely to depend on which promoter(s) they contact, which is inherently related to the 3D structure of the genome [23, 24].

The proximity and frequency of CRE-gene contacts can be measured *in vivo* using chromosome conformation capture (3C) techniques [25]. Chromosome conformation is known to affect how genes are expressed within a cell [26–33]. For example, 3D genome structures may bring linearly distant loci into close proximity, connecting genes with CREs [34–40]. Expressed genes have been observed to spatially localize with distant CREs in 3D FISH experiments [35, 41]. The latest 3C-based technique, Hi-C, pairs the original method’s proximity-based ligation with high-throughput sequencing to identify DNA-DNA contacts on a genome-wide scale [42]. With enough sequencing coverage, Hi-C data can ultimately yield a comprehensive map of the 3D structure of an entire genome at high resolution [43].

It is reasonable to assume that divergence in 3D genome structure can lead to regulatory evolution and ultimately to adaptation of new phenotypes. Currently, however, there are only a few comparative Hi-C data sets that can be used to test this notion, and none in primates. Most Hi-C studies to date have focused primarily on variation in chromatin contact frequencies within a single species, often using transformed or cancerous cell culture lines that may not faithfully represent endogenous phenotypes [44–46]. The few inter-species Hi-C studies we are aware of typically draw comparisons between distantly related species (such as human and mouse), and tend to focus on larger-scale structural variation (i.e. mapping topologically associating domain (TAD) boundaries rather than specific locus-locus contacts) [47–49].

The goal of our study was to comparatively study 3D genome structure and its impact on gene regulation in primates. We collected Hi-C data from iPSCs derived from four human and four chimpanzee individuals to characterize ‘lower-order’ locus-locus contacts and to infer ‘higher-order’ structural features, such as TADs and TAD boundaries. We also quantified gene expression levels using RNA-sequencing data from the same eight cell lines. We overlaid these data with existing functional annotations, including histone marks and chromatin accessibility data, and evaluated the extent to which inter-species variation in 3D genome structure and epigenetic profiles are associated with gene expression divergence between humans and chimpanzees.

## Results

We performed *in situ* Hi-C as previously described [45] on a sex-balanced panel of four human and four chimpanzee integration-free iPSC lines that were previously generated and quality-checked by the Gilad lab [50]. Using HiCUP [51] and HOMER [52] (see Methods), we obtained genome-wide Hi-C contact maps at 10 kb resolution for all eight individuals, with each map containing approximately one billion sequencing reads. Since there is currently no clear gold standard for Hi-C normalization and statistical modeling, we also used an alternative method, Juicer [53], to confirm that our results are robust with respect to other choices of normalization schemes or modeling (Figures S9-14). In the main paper we report the results that were obtained using the HOMER pipeline; results using the alternative pipelines are shown in the supplement in order to document the robustness of our conclusions.

We used HOMER to independently classify about 800,000 significant contacts (p < 0.01) in the Hi-C data obtained from each individual (genomic coordinates of all contacts in all individuals are provided in Tables S1-S8). We define ‘significant contacts’ as a pair of distant, 10 kb regions, that we observed to be in physical proximity more often than expected by chance. Throughout the paper, we refer to Hi-C contacts as ‘lower-order’ or ‘pairwise’ interactions in order to distinguish them from higher-order, chromosome-scale structures (i.e. TADs and TAD boundaries).

Our goal was to perform a comparative analysis of Hi-C contacts in humans and chimpanzees. One intuitive approach to do so might be to identify the orthologous locations of each significant contact in the two species and classify such contacts as shared or unshared. However, this could lead to an inflated estimate of inter-species differences due to incomplete power to identify significant contacts in one species or the other. Instead, we collected the coordinates of all contacts identified as significant in at least one individual into a single database. For each significant contact in the database, we identified the corresponding orthologous regions in the human and chimpanzee genomes and removed non-orthologous regions (see Methods). For each individual, we extracted the normalized contact frequencies between all pairs of loci in the database. We observed that the variance in contact frequency was lower for interactions that were independently identified in a greater number of samples (Figure S1). We thus filtered out interactions that were independently classified as significant in fewer than four individuals, regardless of species. This approach allowed us to compare contact frequencies between species for 347,206 interactions while largely sidestepping the problem of incomplete power.

### Inter-species differences in 3D genomic interactions

The distribution of interaction frequencies across all the genomic contacts we considered are visually similar in the two species (Figure S2), but we nonetheless used pairwise cyclic loess normalization to minimize the effects of technical variables on our data. Principal components analysis (PCA) and unsupervised hierarchical clustering of the Hi-C data revealed that, as expected, samples cluster by species (Figure 1).

**Figure 1.**
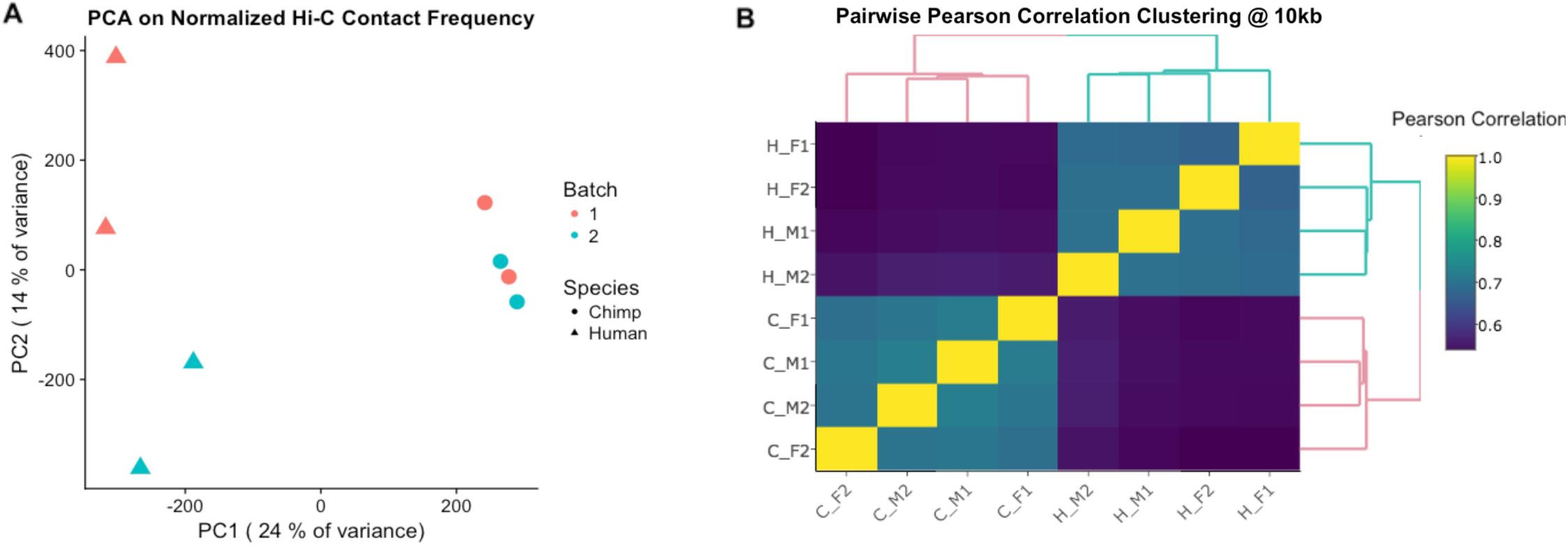
Regulatory landscapes cluster by species. A: Principal components analysis (PCA) of HOMER-normalized interaction frequencies for the union of all significant hits between humans (triangles) and chimpanzees (circles). PC1 is highly correlated with species (r = 0.98 and P < 10^−5^ in regression of PC1 by species). B: Unsupervised hierarchical clustering at 10 kb of the pairwise correlations (Pearson’s r^2^) between HOMER-normalized interaction frequencies. The first letter in the labels demarcates the species (H for human and C for chimpanzee), and the following symbols indicate sex (male, M or female, F) and batch (1 or 2).

To identify inter-species differences in contact frequencies, we analyzed the data using a linear model with fixed effects for species, sex, and processing batch (see Methods). At an FDR of 5%, we classified 13,572 contacts (about 4%) as having differential read counts between humans and chimpanzees. Careful bioinformatic analysis suggested to us that approximately 4,000 of these differences might be explained by errors in assigning orthology (Methods). We conservatively excluded these loci from downstream analyses. We thus classified with confidence 9,629 Hi-C contacts with a significant difference in read counts between the two species. We refer to these contacts as inter-species differentially contacting (DC) regions.

Across all DC regions, 55% exhibited a higher contact frequency in chimpanzees, while 45% showed a higher frequency in humans (Figure 2A). As expected, there is a clear and expected bias towards classification of significant contacts in the species that also shows higher contact read counts (Figure 2A). We observed that some chromosomes were associated with greater asymmetry in inter-species contact frequencies than others (Figure 2B). Most of the chromosomes that exhibited this strong asymmetry are known to have undergone large-scale rearrangements between the human and chimpanzee lineages (e.g. chr. 2 and 17) [54–59].

**Figure 2.**
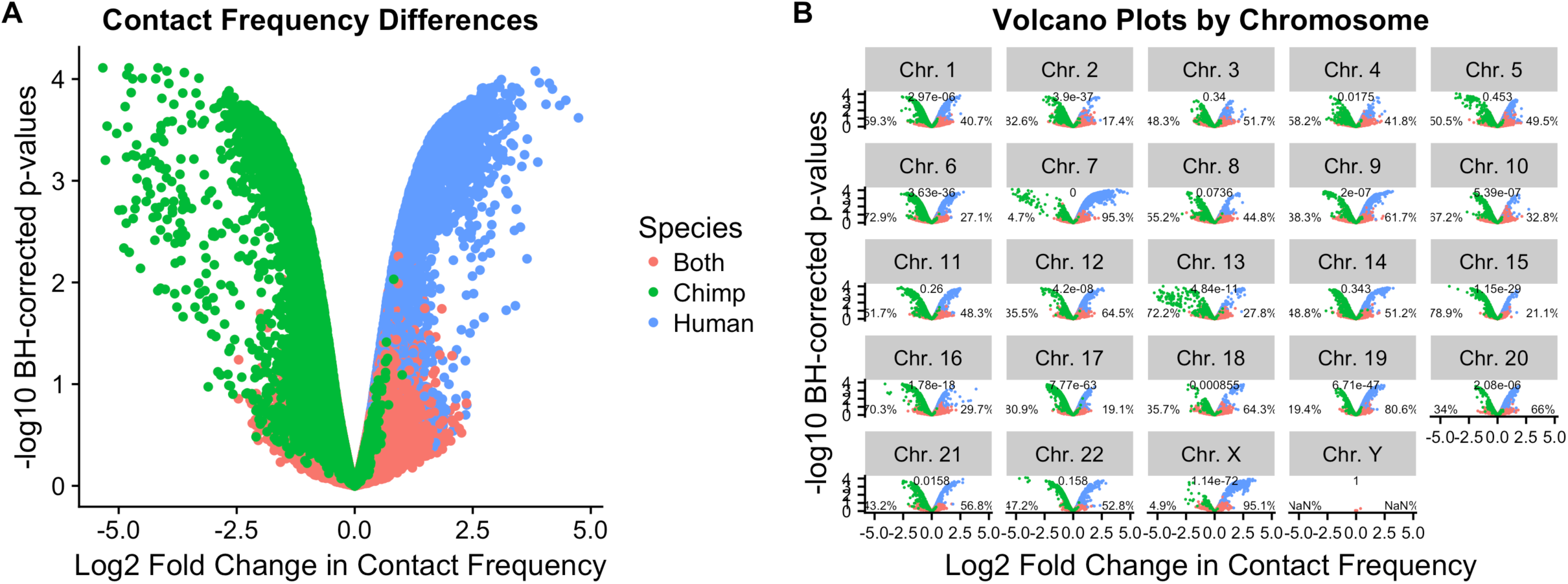
Linear modeling reveals large-scale chromosomal differences in contact frequency. A: **Volcano plot** of log2 fold change in contact frequency between humans and chimpanzees (x-axis) against Benjamini-Hochberg corrected -log10 p-values (y-axis),after filtering out data affected by putative errors in orthology assignment (unfiltered results are plotted in Supplementary Figure 3). Data is colored by the species in which they were discovered as significant. Because humans were coded as a 1 in the linear model, negative values represent pairs with decreased contact frequency in humans compared to chimpanzees, and positive values represent pairs with increased contact frequency in humans. **B: Per-chromosome volcano plot** using the same legend as in A. Numbers above each panel represent p-values from a binomial test under the expectation of significant hits being evenly distributed across the species. Percentages to the left and right of each plot represent the percentage of significant hits that are stronger in chimpanzees and humans, respectively.

Next, we turned our attention to higher-order chromosomal structures by characterizing TADs in each species. To infer steady-state TAD structures, we pooled reads across all individuals within each species to create “high-density consensus” Hi-C maps for humans and chimpanzees [53]. We used the Arrowhead algorithm at 5kb resolution [53] to independently infer TADs and TAD boundaries within species (see Methods). We then used liftOver to identify orthologous genomic regions that corresponded to these TAD boundaries (Supplementary Tables S9-10 list the TAD boundaries identified in each species; Supplementary Table S11 lists the orthologous locations of the combined TAD boundaries). Using this approach, we found that only ~46% of TAD boundaries discovered across humans and chimpanzees are shared between the species (Figure 3). This result was somewhat surprising to us, as the notion in the field is that TAD boundaries are highly conserved even across distantly related species (see Discussion).

**Figure 3.**
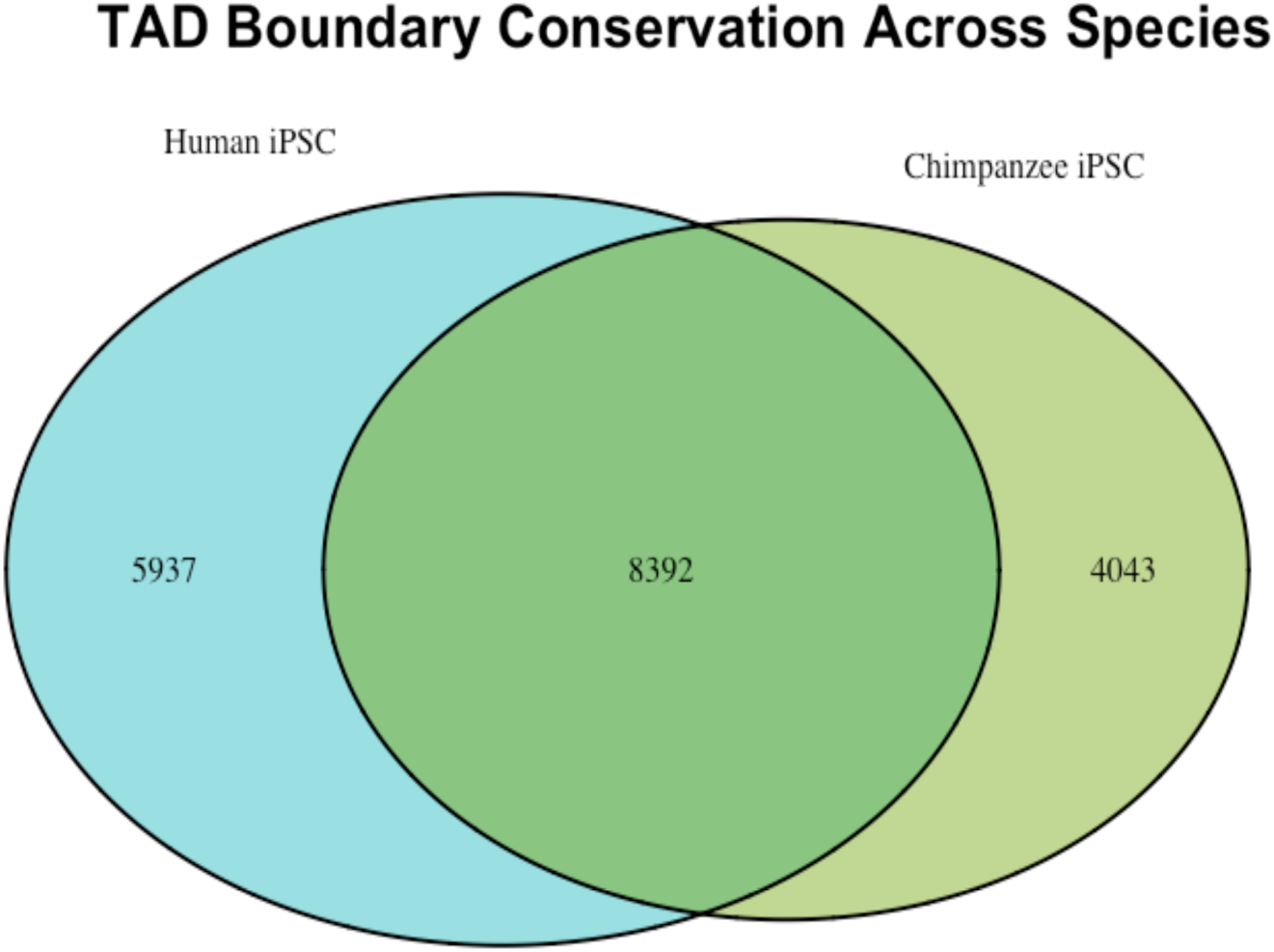
Higher-order chromosomal structure differs substantially between humans and chimpanzees. Venn diagram showing the overlap of TAD boundaries inferred using the Arrowhead algorithm at 5kb on high-density consensus Hi-C maps in each species [53]. Of all TAD boundaries discovered and retained as orthologously mappable in both genomes, ~46% are shared across species. Boundaries found only in humans represent another ~32% of this union list, and boundaries found only in chimpanzees represent the remaining ~22%. We observed remarkably similar estimates of TAD boundary sharing using individual-level data with different algorithms and at different resolutions (Figure S4).

To examine our inference more closely, we also inferred TADs and TAD boundaries on an individual basis, using TopDom [60] with a 200kb window size in each sample (see Methods). We once again used liftOver to identify orthologous genomic regions corresponding to the TADs and TAD boundaries in all individuals (Supplementary Tables S12-S19). Using this approach, about half of the boundaries within each species were identified in only one individual (Figure S4A), and only ~25% of the boundaries were shared between species (Figure S4B). Excluding TAD boundaries that were identified in only one individual, the proportion of shared boundaries between the two species increases to 41, which is in line with the estimate we obtained for the combined data. Our inference of lower than expected conservation of TADs and TAD boundaries is robust with respect to the algorithm used to infer TADs (Figure S4C-D).

### The relationship between inter-species differences in contacts and gene expression

We previously collected RNA sequencing data from the same human and chimpanzee iPSC lines [61]. We jointly analyzed the Hi-C and RNA-sequencing data to learn how often inter-species differences in 3D genomic contact frequencies are associated with inter-species differences in gene expression. We first identified 7,764 orthologous genes for which we have expression and Hi-C data anchored at a region that overlaps the gene’s transcription start site (TSS; see Methods). A single genomic region that overlaps a TSS can have multiple contacts to other genomic regions. For the purpose of our analysis, we conservatively considered only the contact that shows the highest inter-species divergence for each gene.

We did not observe a correlation between gene expression and contact frequency when we considered data from all 7,764 genes. However, when we focused on the 1,537 genes classified as differentially expressed (DE) between humans and chimpanzees (at FDR ≤ 0.05), we observed an excess of both positive and negative correlations between inter-species differences in gene expression and inter-species differences in Hi-C contacts (Figure S5). Indeed, genes whose TSS was associated with inter-species DC were more likely to be DE between species (χ^2^ test; *P* = 0.03; Figure 4A). The association between Hi-C contacts and gene expression divergence was somewhat stronger if instead of focusing on the contact with the highest divergence, we obtained a summary *P*-value [62] for testing the null hypothesis that there are no differences between the species in any of the contacts associated with the TSS for a given gene (*P* = 0.02; Figure 4B).

**Figure 4.**
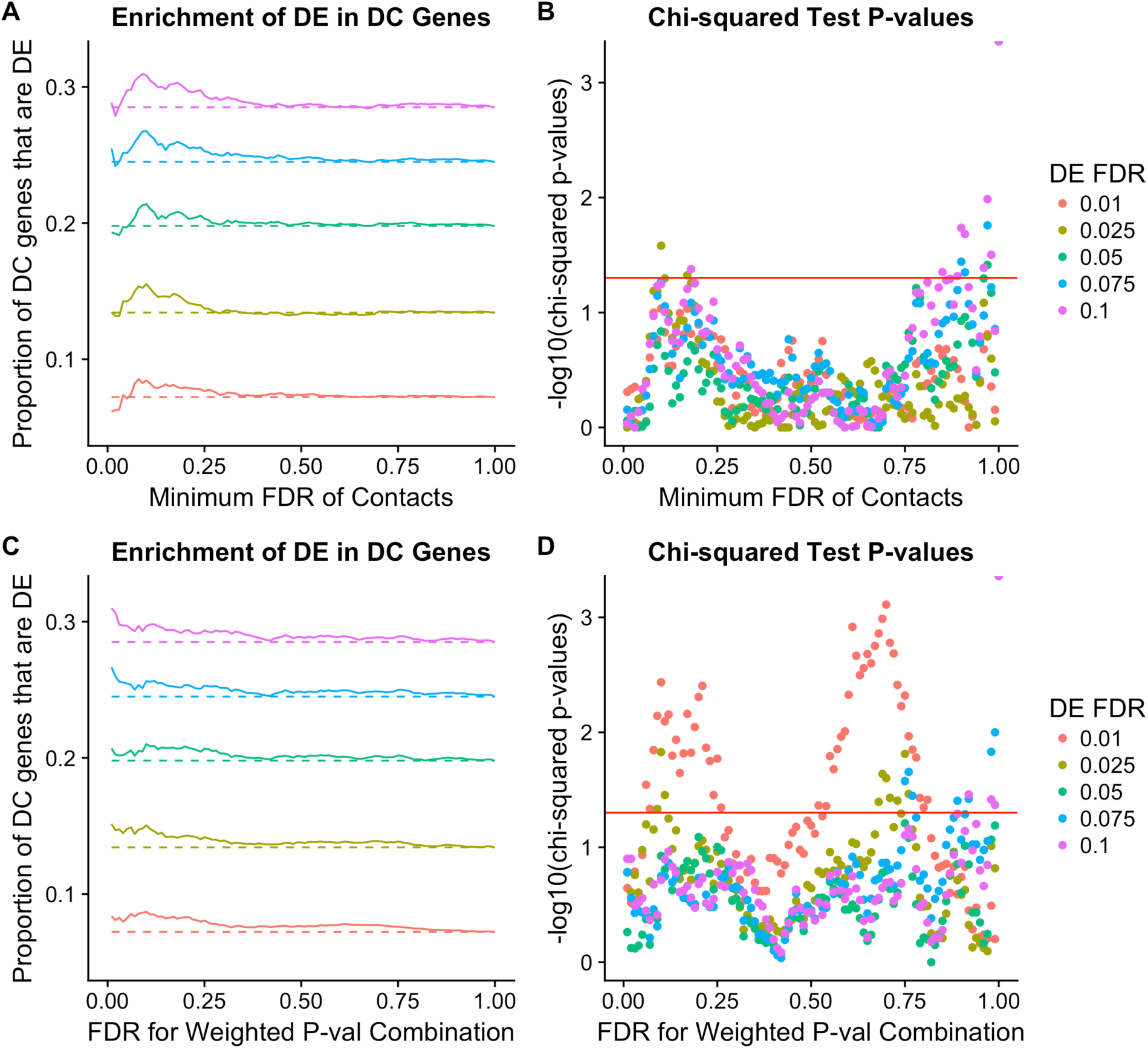
Differentially expressed genes show enrichment for differential Hi-C contacts. **A:** Enrichment of differential expression (DE) in genes with differential Hi-C contact frequencies (DC). The proportion of DC genes that are significantly DE (y-axis) is shown across a range of DC FDRs (x-axis). Colors indicate different DE FDR thresholds and dashed lines indicate the expected proportion of DC genes that would be DE based on conditional probability alone. **B:** Chi-squared significance tests for the enrichments in A. –log10 chi-squared test p-values for enrichment of DC in DE genes (y-axis) is shown for a range of DC FDRs (x-axis). If a gene had multiple DCs, we plotted only the contact that had the minimum FDR. As in A, colors indicate different DE thresholds. A horizontal red line is drawn where p=0.05. The strongest signal of enrichment is observed at 2.5% DE FDR and 10% DC FDR (p=0.03). Because tests are not completely independent, we did not correct for multiple hypothesis testing. **C:** Same as in A, but this time, a weighted p-value combination technique [62] was used to integrate each Hi-C bin’s DC FDR across all of its contacts. **D:** Same as in B, but for the weighted p-value combination instead of the minimum FDR contact.

A combined analysis of functional genomic data does not allow us to infer a direct causal relationship between chromatin contacts and gene expression patterns. Nevertheless, independent evidence strongly suggests that changes in 3D genomic structure can affect interactions between regulatory elements and promoters [45, 63-66], which may ultimately drive differences in gene expression levels [33, 63-68]. We thus sought to quantitatively estimate the extent to which inter-species DC might explain gene expression differences between the species. We performed a mediation analysis by using linear models to estimate gene expression divergence before and after accounting for the corresponding contact frequencies (see Methods). Specifically, we applied an empirical Bayes adaptive shrinkage approach [69] to compare the gene expression divergence effect size estimated under each model. Approximately 19% of DE genes (294/1537) had a smaller divergence effect size (at s-value ≤ 0.05) after accounting for their corresponding inter-species DC data (Figure 5A). As a control, we repeated this analysis for genes that were *not* classified as DE between species and, as expected, we found no effect of contact frequency on divergence effect size (Figure 5B). Taken together, these data suggest that inter-species variation in gene expression levels can often be explained by divergence in Hi-C contacts. Such a model is supported not only on the basis of our genome-wide data analyses, but also when observing a single integrative example, examining differential contacts as well as differential expression across the species (Figure 6).

**Figure 5:**
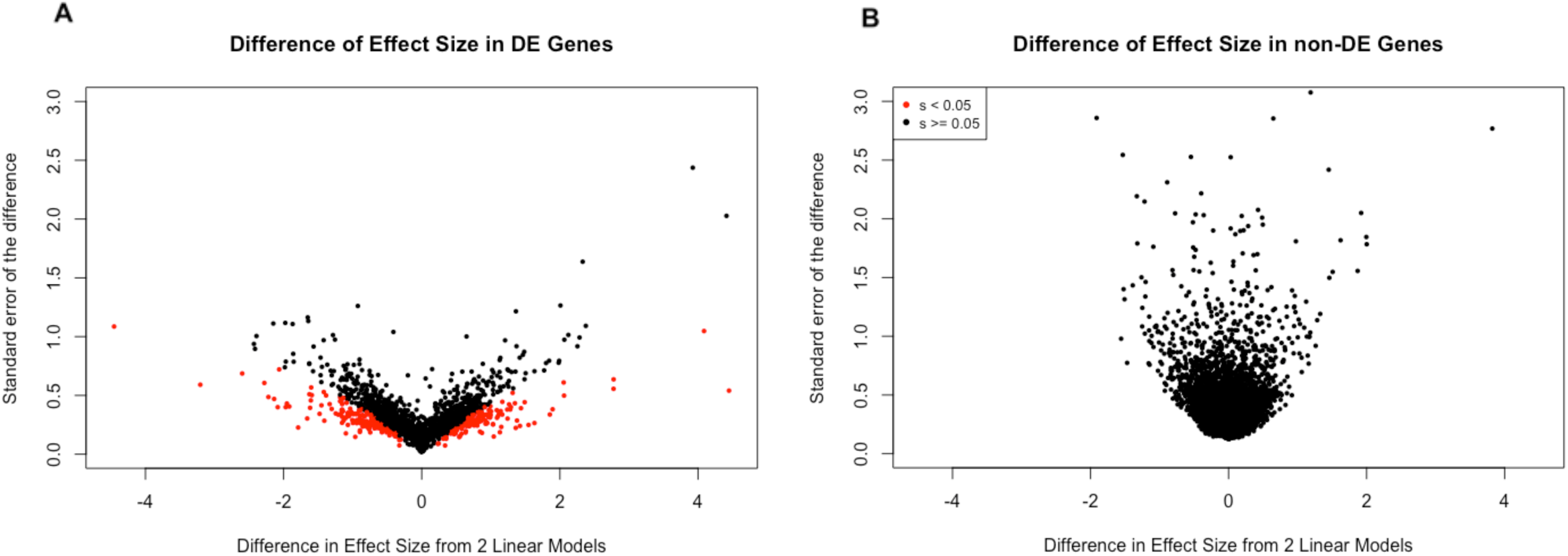
Gene expression variance is explained by chromatin contacts for 19% of DE genes. **A:** Volcano plot of the differences in DE effect size between models before and after conditioning on contact frequency (x-axis) plotted against the standard error of the difference (y-axis). Adaptive shrinkage was used to evaluate significance of the differences in effect size. Amongst DE genes, 19% (294/1537) of genes showed a statistically significant and sign-confident effect of Hi-C contacts on expression levels (s-value ≤ 0.05). **B:** Same as in A, but performed on the set of 6,227 non-DE genes. No genes showed a statistically significant difference in effect size between the two models.

**Figure 6:**
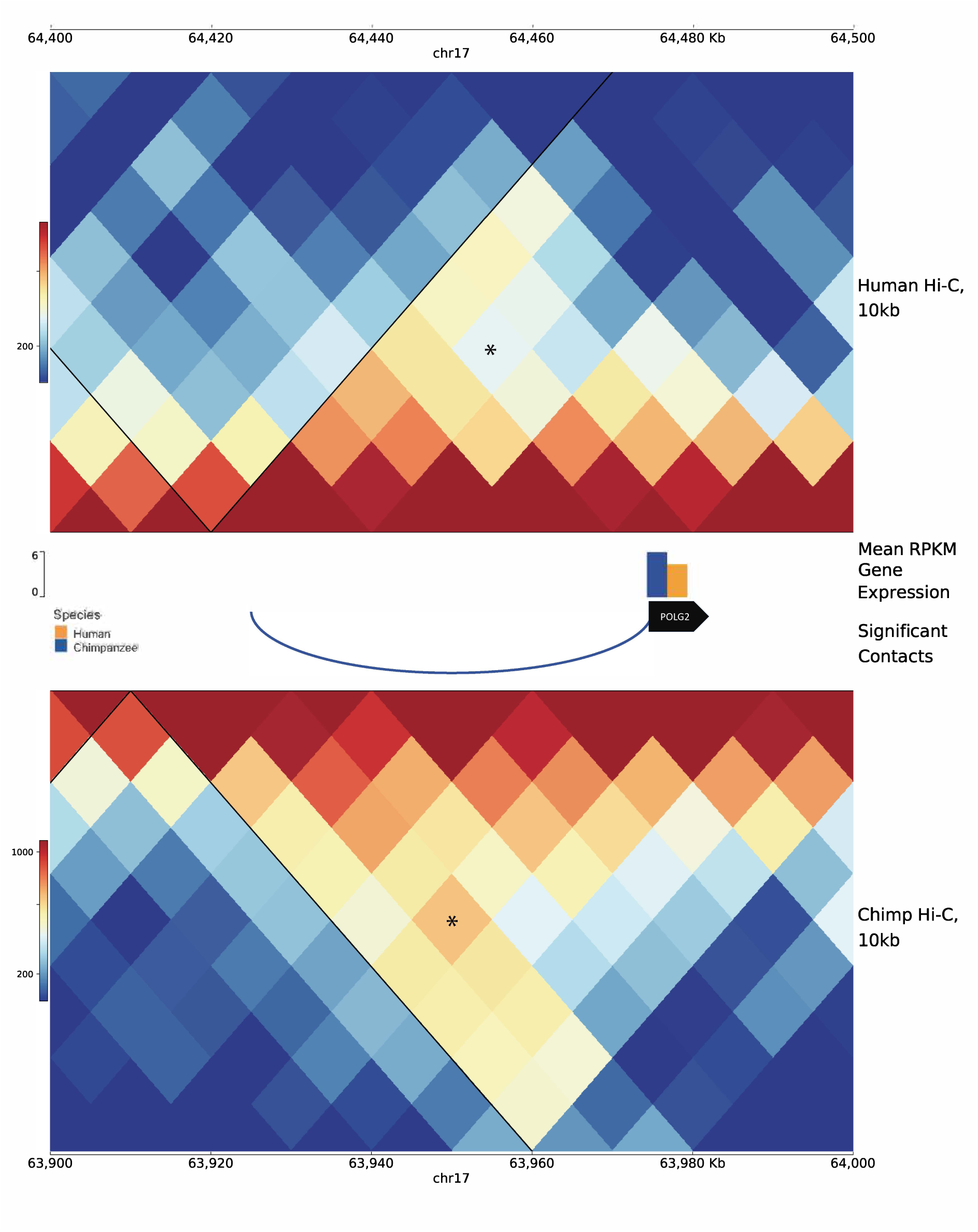
A differentially expressed gene mediated by differential contact. PyGenomeTracks plot [70] of a 100kb syntenic region between species, with a representative example of a differentially expressed gene influenced by differential contact (s=0.04). Top and bottom of plot show Hi-C matrices at 10kb resolution for humans and chimpanzees, respectively, with TADs overlaid as black lines. In center, beneath human Hi-C, is a track displaying mean RPKM gene expression for the gene POLG2 within each species (humans in orange and chimpanzees in blue). Underneath expression values is an arc diagram displaying a differential contact observed to be stronger in chimpanzees in our dataset. Asterisks on the Hi-C matrices indicate the specific orthologous locus pairs that were tested for differential contact between species. This contact was also discovered as significant in HOMER in all four chimpanzee individuals, whereas HOMER did not infer any significant human contacts within this region. A broader view of this same region is presented in Supplementary Figure 15.

### The chromatin and epigenetic context of inter-species differences in 3D genome structure

Finally, we reasoned that species-specific contacts would be more likely to involve active, functional regulatory elements. To test this hypothesis, we assessed the overlap between our Hi-C data and publicly available chromHMM annotations based on histone modification data from human embryonic stem cells [18]. We assigned each Hi-C locus to an epigenetic state based on its maximum weighted base pair overlap with 15-state chromHMM annotations (see Methods and Figure S6). Our approach to classify Hi-C regions with a functional assignment based on majority sequence overlap is arbitrary, but our conclusions are robust with respect to alternative approaches (Figures S6 and S13).

We found marked differences in the chromHMM annotations between genomic regions that are inferred to physically contact a promoter and those that do not contact a promoter (Figure 7A). For example, genomic regions in physical contact with a promoter are enriched with genic enhancer annotations (χ^2^ test; *P* = 0.002), as might be expected. Perhaps more novel is the observation that inter-species DC regions were also enriched with genic enhancers, in contrast to regions that did not differ in contact frequency between the two species (*P* = 0.04; Figure 7E). We note that this latter observation is not robust with respect to different annotations of enhancers, and we do not find this association if we simply combine all regions annotated as ‘enhancers’ in the data set.

**Figure 7.**
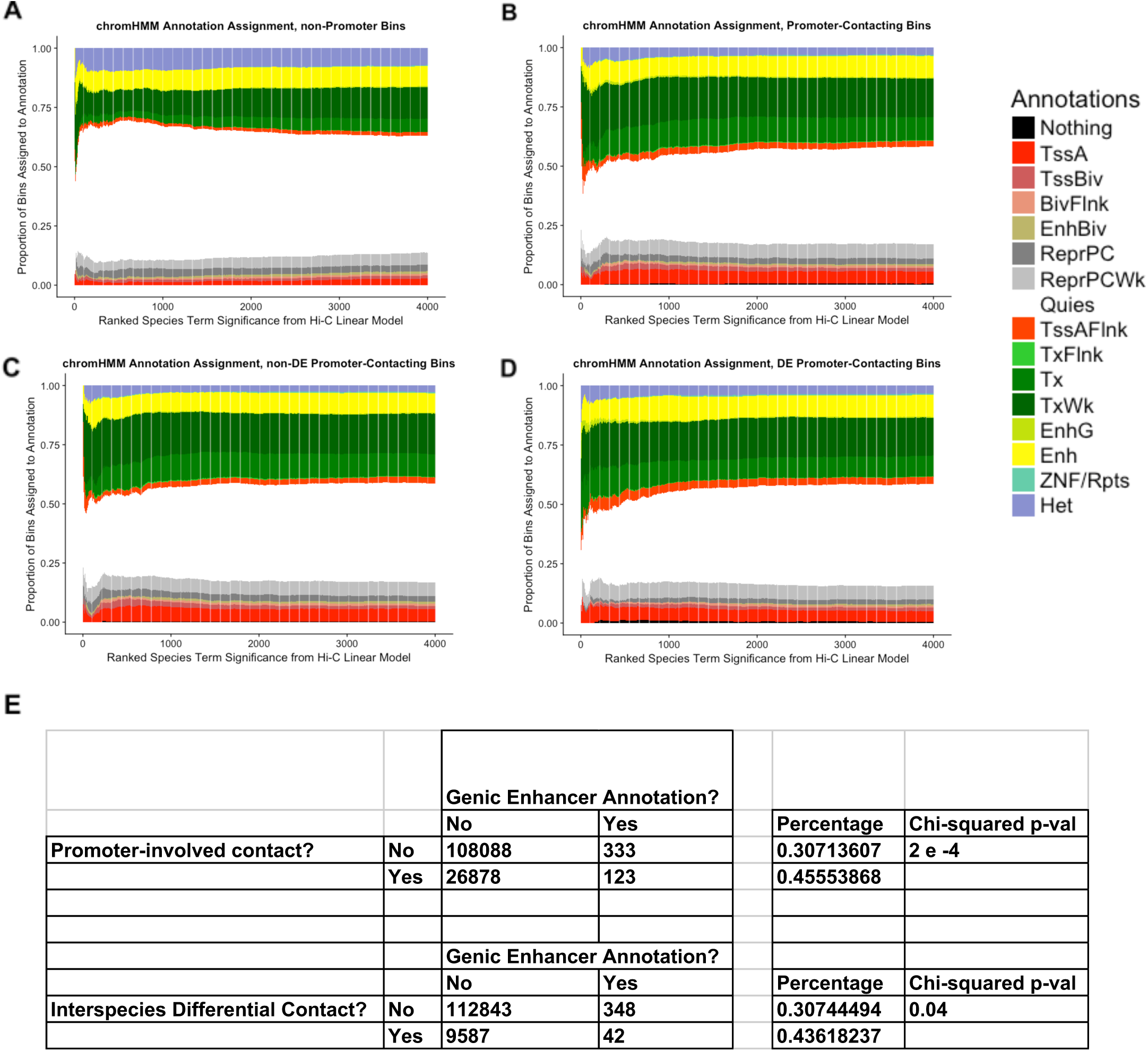
Dynamics of chromHMM state among significant Hi-C contacts. In A-D, Hi-C contact loci are ranked in order of decreasing significance (x-axis), with species-specific contacts (i.e. lower FDR for our species term from linear modeling) on the left, decreasing in significance towards the right. The y-axis shows cumulative proportion of chromHMM annotation assignments for all Hi-C loci at the give rank or lower. (TssA-Active TSS, TSSBiv-Bivalent/Poised TSS, BivFlnk-Flanking Bivalent TSS/Enh, EnhBiv-Bivalent Enhancer, ReprPC-Repressed PolyComb, ReprPCWk-Weak Repressed PolyComb, Quies-Quiescent/Low, TssAFlnk-Flanking Active TSS, TxFlnk-Transcription at gene 5’ and 3’, Tx-Strong transcription, TxWk-Weak transcription, EnhG-Genic Enhancers, Enh-Enhancers, ZNF/Rpts-ZNF genes and repeats, Het-Heterochromatin) **A:** chromHMM annotation assignment for non-promoter contacting bins. **B:** chromHMM annotation assignment for all promoter-contacting bins. **C:** chromHMM annotation assignment for non-DE promoter-contacting bins. **D:** chromHMM annotation assignment for DE promoter-contacting bins. **E:** Two-by-two tables of the percentage of contacts that we assigned to the “genic enhancer” annotation (see Methods). Contacts are separated out on the left by whether they involved a promoter in the top table, and whether they showed evidence for inter-species DC on the bottom table. P-values from a chi-squared test are shown on the right.

We repeated the enrichment analysis of Hi-C regions using existing human iPSC histone mark data, including H3K27ac, H3K4me1, and H3K4me3, H3K27me3 and DNase I hypersensitivity site (DHS) data collected in h1-hESCs [18]. As expected, Hi-C regions in contact with a promoter showed greater overlap with DHS peaks than Hi-C regions that did not contact a promoter (t-test, *P* < 10^−15^; Figure S7A). When we focused on contacts involving a promoter, we found that inter-species DCs that are also associated with DE genes showed the largest overlap with DHS peaks, followed by DC regions that were not associated with DE genes (*P* = 0.002). Regions that were not associated with either DC or DE showed the least amount of overlap with DHS (*P* = 0.0001; Figure S7B).

Remarkably, apart from the heterochromatic, repressive marker H3K27me3 (where the sign of the effect was the same, but the enrichment was not significant), Hi-C regions that are DC and are also associated with DE genes are more likely to overlap all other histone marks in our data set compared with Hi-C regions that are not DC and are not associated with a DE gene (all enrichment *P* < 0.04; Figure 8A-D). In other words, inter-species DCs associated with DE genes are more likely to occur in genomic regions that are marked by histone modification, and are thus likely to have a regulatory function.

**Figure 8.**
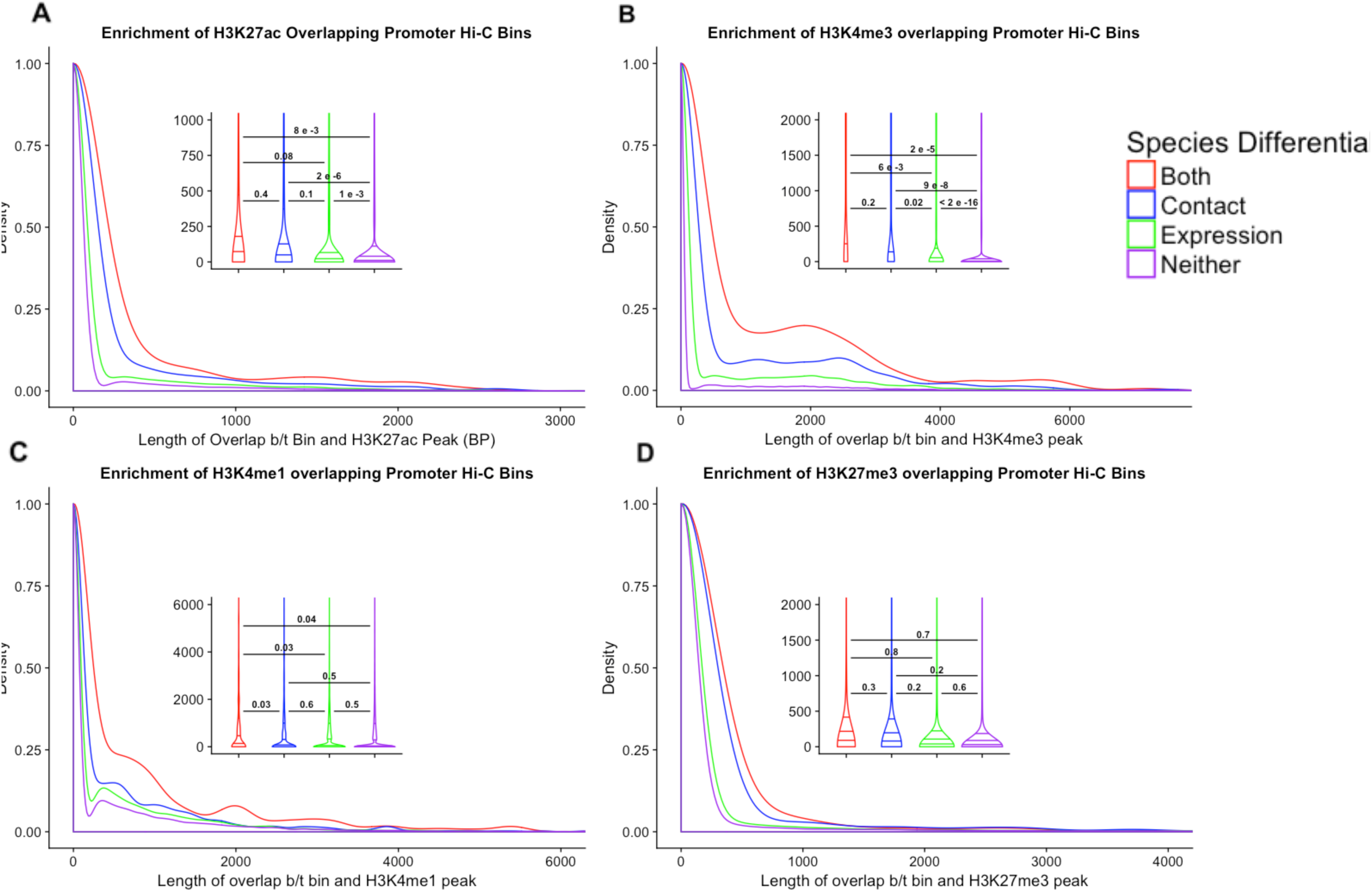
Overlap of activating and repressive histone marks among Hi-C contacts. A-D show density plots of the base pair overlap between different classes of Hi-C contact loci and various histone marks. All histone mark data were obtained from ENCODE in experiments carried out in human iPSCs (A-C) or hESCs (D). We grouped contacts into 4 classes, indicated by color: those that show differential contact between species, those that show differential expression between species, those that show both, and those that show neither. We used pairwise t-tests to compare differences in the mean overlap among the four classes of Hi-C loci. Inlaid violin plots show the significance level of each t-test, with horizonal lines connecting the two groups used for comparison. Lines inside each violin represent the 25^th^, 50^th^, and 75^th^ percentiles of the data. **A:** Density plot of base pair overlap between Hi-C contact loci and the active histone mark H3K27ac. **B:** Same as in A, but for the active histone mark H3K4me3. **C:** Same as in A and B, but for the active histone mark H3K4me1. **D:** Same as in A-C, but for the repressive histone mark H3K27me3.

## Discussion

In general, we observed lower-order, pairwise chromatin contacts in iPSCs to be conserved between humans and chimpanzees (only about 4% of pairwise contacts were classified as quantitatively different between the species). We believe that this observation is intuitive, though we acknowledge that with only four individuals from each species, we were underpowered to identify small, quantitative inter-species differences in contact frequencies.

In contrast to the conservation of lower order pairwise contacts, we did not find higher-order chromatin structures, such as TAD boundaries, to be generally conserved between human and chimpanzee iPSCs. This observation seems to contradict previous reports suggesting that TADs are strongly conserved across tissues and species [44, 45, 71-74]. In particular, Dixon *et al.* [71], who originally described TADs, reported that TAD boundaries were conserved in human and mouse embryonic stem cells. However, this comparison was made using lower-resolution Hi-C data (40kb), and with a somewhat relaxed definition of orthology to determine synteny. Rudan *et al.* [44] reported general conservation of TAD boundaries in liver cells from rhesus macaque, dog, rabbit, and mouse, but did not report the number or proportion of conserved regions they observed. Moreover, this study did not perform a genome-wide assessment of TAD boundary conservation across species. This is not unusual; several other comparative studies reporting that TAD boundaries are highly conserved did not perform a genome-wide analysis [44, 45, 72-74]. Rather, these studies observed conservation among certain characteristics of the boundaries, such as enrichment for CTCF motifs in a convergent orientation, or enrichment for other epigenetic marks.

Based on our analysis of the literature, we believe that the common notion that TADs are highly conserved in their placement across species is not well supported. Indeed, recent evidence from different *Drosophila* tissues [75] and plant species [49] suggests that TADs and TAD-like domains may not be particularly conserved, which raises questions about the stability of these higher-order structures and the significance of their role in the evolution of gene regulation across different lineages. However, the extent (or lack) of inter-species TAD conservation is difficult to falsify with existing data, partially because there is no standard method for identifying TADs and comparing them across species. The ability to reliably identify TADs also depends on the quality of the genome assemblies used, the approach for inferring synteny, sequencing depth and coverage, and various other parameters.

In our case, we considered multiple strategies to define and analyze pairwise chromatin interactions and higher-order structures in humans and chimpanzees, using a conservative approach to identify syntenic regions. Our observation that TAD boundaries are not highly conserved between humans and chimpanzees is robust across the various methods we tested. Thus, we are confident that our data do not support a model in which large-scale domains remain invariant across evolutionary time.

### Contribution of Variation in 3D Genome Structure to Expression Divergence

We considered our Hi-C data along with gene expression data previously collected from the same cell lines [61] and assessed the extent to which inter-species variation in 3D genome contacts could potentially explain gene expression divergence between species. Previous studies have observed that spatial co-expression of genes is associated with chromatin interaction profiles [26, 30, 31, 41, 76]. A number of studies have focused on differentially expressed genes following a treatment or perturbation and observed that such genes are often associated with corresponding differences in nearby chromatin contacts [63, 64]. Consistent with these reports, we found an enrichment of inter-species differences in pairwise chromatin contacts that involve promoters of differentially expressed genes between the species. Our observations are robust with respect to a range of data processing decisions and the statistical cutoffs we used. Under the common assumption that changes in chromatin contacts are more likely to explain differences in gene expression than vice versa, our results support the notion that species-specific 3D genomic contacts play an important role in the evolution of gene regulation.

Our observation that inter-species differences in pairwise genomic contacts are associated with regulatory evolution more than differences in large scale TAD boundaries is also consistent with previous reports. For example, Rao *et al*. [33] found that the degradation of cohesin, one of the proteins involved in maintaining TAD boundaries and large-scale loops, is associated with only modest effects on gene expression. In contrast, a number of other studies found strong correlations between differences in fine-scale genomic contacts and differences in the expression of nearby genes [45, 65, 66].

Previous studies have identified a wide variety of regulatory phenotypes that contribute to inter-primate differences in gene expression levels [10, 11, 13, 77-80]; 3D genome conformation is only one of the putative upstream factors in the evolution of gene regulation. Our results argue for a model whereby inter-species differences in pairwise contact frequencies are among the main drivers of expression divergence between humans and chimpanzees. Given the low 10-kb resolution of our Hi-C data, it is likely that we have underestimated the contribution of inter-species variation in 3D genome structure to gene expression divergence between species. Future comparative Hi-C studies that sequence deeply enough to obtain higher, sub-kilobase resolutions, will allow researchers to resolve variation in contact frequency at even smaller scales, augmenting predictive power.

### Functional Annotations

Finally, we considered our data in the context of functional chromatin annotations available for the human genome. Previous studies have shown that 3D contact maps produced by Hi-C can be accurately recapitulated by epigenetic marks [81, 82]. Other reports have found enrichments for various chromatin accessibility and histone marks among interactions inferred from chromosome conformation capture data [83, 84].

Our results corroborate and expand upon these findings. The differences we observed in chromHMM state assignments in our comparisons (namely, more active and less repressive states in promoter-involved contacts and contacts overlapping differentially expressed genes), provide additional support for the functional relevance of our inferences. Indeed, while our study design does not allow us to directly infer causality between chromatin interactions and gene expression, the functional enrichments we observed for different epigenetic marks suggest that 3D genome conformation may be one of the upstream elements in the chain of events driving the evolution of gene expression. Although this notion is intuitive to us and is consistent with our data, it is still possible that differences in epigenetic marks are the true drivers of divergence in gene expression levels and/or chromatin contacts between humans and chimpanzees.

Future studies integrating similar data types could explore these possibilities by examining epigenetic marks across species (only human data were available to us), which would enable researchers to polarize the regulatory differences in orthologous sequences between humans and chimpanzees. This would also allow for a sharper definition of the functional classes of inter-species differences in lower-order chromatin contacts.

## Materials and Methods

### Induced Pluripotent Stem Cells (iPSCs)

As described previously, the Gilad lab has derived panels of both human and chimpanzee iPSCs via episomal reprogramming [50]. We collected human fibroblasts with informed consent obtained from all human participants under University of Chicago IRB protocol 11-0524,. We obtained fibroblasts from chimpanzees from the Yerkes Primate Research Center of Emory University under protocol 006-12, in full compliance with IACUC protocols [50]. All experimental methods are in accordance with the Helsinki Declaration. To ensure their quality, we validated iPSCs from both species as pluripotent at high passages (>10). Quality control checks included an embryoid body assay confirming their ability to differentiate into all three germ layers, qPCR of endogenous transcription factors associated with pluripotency, PCR to confirm the absence of exogenous pluripotency genes (both from residual episomal plasmid or genomic integration), and PluriTest [85], a bioinformatics classifier that assesses pluripotency based on gene expression data. In the current study, we grew all cell lines in the same incubator in two passage-matched batches, which were also balanced across species and sex, in order to avoid batch effects.

### In-situ Hi-C Library Preparation and Sequencing

We performed *in situ* Hi-C with the restriction enzyme MboI, as previously described

[45] on the iPSCs from both species. This version of Hi-C uses intact nuclei, which confers the advantage of reducing the frequency of spurious DNA-DNA contacts that result from random proximity ligation in solution. We grew cells in feeder-free conditions [86] to approximately 80% confluence before adding formaldehyde to crosslink the proteins mediating DNA-DNA contacts. We flash-froze pellets of 5 million cells each before beginning the *in situ* Hi-C protocol [45]. We used MboI to cut the DNA at each of its 4-bp recognition sites (GATC) throughout the genome. Ligation of proximal fragments with T4 DNA ligase yielded chimeric DNA molecules representing two distinct loci. Libraries were created in two balanced batches identical to the cell growth batches and sequenced (100bp paired-end) on an Illumina Hi-Seq 4000 at the University of Chicago Genomics Core Facility. In order to avoid batch effects resulting from differences in flow cells, libraries were sequenced across three lanes, each on separate flow cells balanced for species.

### Hi-C Read Mapping, Filtering, and Normalization

We preprocessed, mapped, and filtered the resulting FastQ sequence files using HiCUP [51]. We also used HiCUP to truncate the reads at ligation junctions. Thereafter, we usedbowtie2 [87] to independently map the two mates of paired-end sequences to either the hg38 or panTro5 genomes, and removed reads with MAPQ < 30. We carried out further HiCUP filtering as described based on an *in silico* genome digest in order to remove experimental artifacts [51]. We then used HOMER, a foundational statistical analysis suite for Hi-C data [52], to tile the genome into a matrix of 10kb bins and assign reads to their corresponding intersecting bins. We subsequently used HOMER to normalize Hi-C contact bins as previously described [52], accounting for known technical biases in Hi-C data. Finally, we called statistically significant interactions independently in each individual using HOMER, based on a null expectation of read counts falling into bins in a cumulative binomial distribution [52]. We retained interactions with a *P* value ≤ 0.01; this is the default recommendation by HOMER. As other studies have noted [53], a traditional multiple testing correction paradigm is overly conservative for Hi-C data due to the high number of tests, and because the spatial nature of the data means that individual tests are highly correlated (and thus not independent).

### Creation of a Union List of Orthologous Hi-C Contacts Across Species

In order to ensure that the contact frequencies we compared across species were from representative orthologous sequences in humans and chimpanzees, we used liftOver with a reciprocal best hits method [88, 89] to transfer interaction bin coordinates across genomes. For all contacts in this union list, we then extracted the HOMER-normalized interaction frequencies from each individual’s 10 kb Hi-C matrix. Including interactions discovered in fewer than 4 individuals increased the variance in our data (Figure S2). Therefore, we retained only the Hi-C contacts that were independently discovered by HOMER in at least 4 individuals, for a total of 347,206 interactions.

### Linear Modeling of Hi-C Interaction Frequencies

In an effort to quantify inter-species differences in the Hi-C interaction frequency values, we parameterized a linear model:

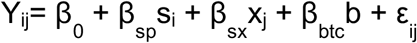

Y_ij_ represents the Hi-C interaction frequency of individual j and species i. β_0_ is the intercept. β_sp_, β_sx_, and β_btc_ are effect sizes for species, sex, and batch, respectively, with their classifier categorical variables s_i_, x_j_, and b, and an error term ε_ij_. We used the R package limma [90] to run significance tests on this model, resulting in 54,273 Hi-C contact pairs where the species term is significant *(P* ≤ 0.05). We applied Benjamini-Hochberg multiple testing correction to find 13,572 interaction pairs significant at 5% FDR.

Initial visualization of the linear modeling results for the species term revealed a stark asymmetry (Figure S3A) suggesting that on a global level, the contacts identified as significant in chimpanzees were much stronger than those identified in humans. Since this did not make biologically meaningful sense, we reasoned that this asymmetry could be due to a technical factor. For example, liftOver conversion of genome coordinates between species can create differences in both the Hi-C locus size and in the genomic distance between mates of a contact pair. We investigated the impact of these two factors on the proportion of contacts classified as differential across the species using a linear model. We discovered that while changes in Hi-C locus size had little effect on the proportion of interspecies DCs, liftOver-induced changes in mate pair distances > 20 kb created a noticeable inflation in this proportion at an FDR of 5% (Figure S3B). We note that it is impossible to ascertain the relative biological and/or technical relevance of the differences seen in these contacts. We thus took a conservative approach to minimize false positives and removed contacts with a >20kb distance difference between species from our downstream analyses (Figure S3C), accepting that some of the inter-species differences we observe may be underestimated.

### Identification of Orthologous Topologically Associating Domains (TADs) and Boundaries

We chose to perform TAD analyses on both individual-level data and on representative species “consensus” data. For our analysis comparing TAD boundaries on species “consensus” Hi-C maps, we combined all the preprocessed Juicer files from all our individuals within a species and used the *juicer_mega.sh* script [53] to create higher density contact maps for each species. We then ran the Arrowhead algorithm at 5 kb resolution to infer TADs, and then we extended the edges of TADs 7.5 kb in each direction to create 15 kb boundaries (accounting for imprecision in boundary inference). We used the reciprocal best hits liftOver strategy to obtain a union list of all orthologously mappable boundaries, and defined a final union set of TAD boundaries using the *intersect* and *merge* functions in bedtools [91]. We estimated the percent of conserved TADs between species by dividing the number of boundaries found in both species (“shared”) by the total number of unique boundaries in the final union list.

To examine individual-level data, we used TopDom [60] with a 200 kb window size to call TADs independently in each individual sample. To obtain robust sets of boundaries found in at least two, three, or four individuals from within the same species, we performed pairwise bedtools [91] overlaps and merges on the boundary elements output by TopDom. Just as we did for the Hi-C contacts, we used the reciprocal best hits liftOver method to obtain a set of orthologous domains and boundaries. We then used these data to obtain a second estimate of TAD conservation between species. We calculated the overlap of orthologous boundaries found independently in both species (“shared”) and divided this number by the total number of unique boundaries in the final union list. We followed the exact same paradigm when analyzing individual-level TAD inferences we produced with the Arrowhead algorithm [53] at 10kb resolution.

### Differential Expression Analysis

Previously, the Gilad lab generated RNA-sequencing expression data on the same iPSC lines from this study (GEO accession GSE110471) [61]. Using these data, we filtered out lowly expressed genes based on a log2 RPKM normalization cutoff of 0.4 that produced a similar proportion of retained genes when using a log2 CPM normalization cutoff of 1.5 (11,946 genes versus 11,292 genes, respectively). We retained only genes where at least half (2) of the individuals in each species showed log2 RPKM ≥ 0.4. This filtering process retained 11,946 genes, which we analyzed with limma [90] using a simple linear model that only included a species term. Using this method, we found 2,268 differentially expressed genes (at 5% FDR).

### Broad Integration of Hi-C and Gene Expression Data

We obtained the overlap between our gene expression data and our Hi-C data by applying bedtools *overlap* [91] to the Hi-C loci and the first exon of each gene. Using a curated file of orthologous gene coordinates between humans and chimpanzees [61], we extracted a one-base-pair interval at the beginning of each first exon to use as a proxy for transcription start sites (TSSs). Given the large size of our Hi-C loci (10 kb), we anticipated that the overlapping contacts would be representative of the genes’ TSSs. Nonetheless, we checked where our one-base-pair intervals fell along the length of the Hi-C loci and confirmed that the data followed a relatively uniform distribution (not shown).

As described in the main text, the difference in dimensionality between the two datasets presented a challenge. While every gene has only one expression value per individual, a given Hi-C locus can and frequently does make contact with many other loci. When a given gene overlapped a Hi-C locus making multiple contacts, we chose the contact with the lowest species term FDR from our linear model (i.e. the most species-specific contact) to represent the interaction frequency for that gene. Accordingly, we interpreted the FDR-adjusted *P* value for the chosen contact as the gene’s differential contact significance.

### Enrichment of Differential Expression in Differential Contacts

We cumulatively tabulated the proportion of genes that had both differential contact and differential expression across a continuous range of differential contact cutoffs and a discrete range of differential expression FDR cutoffs (1%, 2.5%, 5%, 7.5%, and 10%). We then compared these proportions at each FDR to those that would be expected based on conditional probability alone (dotted lines in Figure 4). Finally, we utilized Pearson’s chi-squared test to assess the statistical significance of the enrichment at each FDR (Bottom panel of Figure 4).

### Assessing the Quantitative Contribution of Hi-C Contact Frequencies to Gene Expression Levels

In order to quantify the effect of Hi-C contact interaction frequencies on RPKM gene expression values, we parameterized and compared two separate linear models [79]. The first model is identical to the one described above for the RNA-sequencing data: a simple linear model attempting to predict the expression level of each gene using only a species term. The second model was completed in two steps. First, we predicted the expression level of each gene using its corresponding Hi-C interaction frequency as the only predictor variable. We interpreted the residuals from this analysis as expression values “corrected” for Hi-C interaction frequencies. Next, we input the residual values into the same initial differential expression model as before (with species as the only term).

For each gene, we computed the difference in effect sizes between the two models (one run on RPKM expression values and one run on “Hi-C corrected” expression values), as well as the standard error of the difference. We then used an empirical Bayes approach to shrink both the variance and the effect sizes with the R packages *vash* and *ash*, respectively [69, 92]. We further utilized *ash* to test all genes for significant differences in their effect sizes between the models. Rather than using q-values, however, we called genes’ expression statistically significantly affected by Hi-C contacts if they had an s-value (false sign rate) ≤ 0.05 [69]. Using s-values rather than q-values not only takes significance into account but has the added benefit of assessing our confidence in the direction of the effect.

### Integration with Epigenetic Annotations

We obtained chromHMM 15-state model peak calls in human iPS-18C cells from ENCODE [18] (Table S20). We subsequently found the overlap between the human coordinates of our orthologous Hi-C contact loci and the chromHMM peak calls and quantified the extent of base pair overlap between each locus and all overlapping chromHMM peaks. Because we wanted to speak broadly to epigenetic profile dynamics of our Hi-C loci, we sought to assign each individual locus a single chromHMM annotation based on whichever peak had the highest base pair overlap with that locus. However, the distribution of overlaps of different chromHMM annotation peaks with our Hi-C bins were quite variable in size. To account for this, we normalized each annotation’s overlap length in each locus by multiplying it by the reciprocal of its mean base pair overlap across all our bins (Figure S6). After removing duplicate Hi-C loci, we then assigned individual loci to chromHMM annotations based on these normalized base pair overlaps. To characterize epigenetic profiles of our Hi-C loci, we then took the top ten most differentially contacting loci (i.e. the ten lowest FDR loci from our Hi-C linear modeling), and tabulated proportions of which annotations were represented amongst them. We then iteratively added the next-lowest FDR contact (i.e. two Hi-C loci at a time) to this tabulation, re-calculating proportions on the new set of contacts. We ran this same cumulative proportions analysis separately on contacts not overlapping promoters, contacts overlapping promoters, contacts overlapping promoters of DE genes, and contacts overlapping promoters of genes that were not DE (Figure 7).

We also obtained data on H3K4me1, H3K4me3, and H3K27ac collected in human iPS-18A cells, and data on H3K27me3 and DNase hypersensitivity sites collected in H1-hESCs, all from ENCODE [18] (Table S20). We used bedtools *intersect* [91] to find the base pair overlap between each of these different marks and our Hi-C contact loci. We then removed duplicate Hi-C loci from the dataset and used a pairwise t-test to identify significant differences in the overlapping distributions for different sets of Hi-C classes (based on differential contact and differential expression, Figure 8).

## Supporting information

## Data Availability

The novel Hi-C data collected in this study is available upon request and will be released through GEO upon publication. All analyses performed in R, including creation of most of the figures, can be found at https://ittaieres.github.io/HiC/

## Authors’ Contributions

YG and IEE conceived of the study and designed the experiments. IEE performed the experiments. IEE and KL analyzed the data, with input and code from CJH and LEB. IEE and YG wrote the paper. All authors read and approved the final manuscript.

## Acknowledgement

We thank the members of the Gilad, Nobrega, and Stephens labs for helpful discussions, particularly Matthew Stephens, Bryan J. Pavlovic, Débora R. Sobreira, and Lindsey E. Montefiori. We thank Natalia Gonzales for help editing the paper. This work was supported by NIH grant GM122930 and NIH grant HG002585. IEE was supported by the Genetics and Gene Regulation Training Grant (T32 GM07197). LEB was supported by the National Science Foundation Graduate Research Fellowship (DGE-1144082) and by the Genetics and Gene Regulation Training Grant (T32 GM07197). The content presented in this article is solely the responsibility of the authors and the funding bodies had no role in the design of the study nor in collection, analysis, interpretation of data, and writing the manuscript.

---

**Figure S1.**
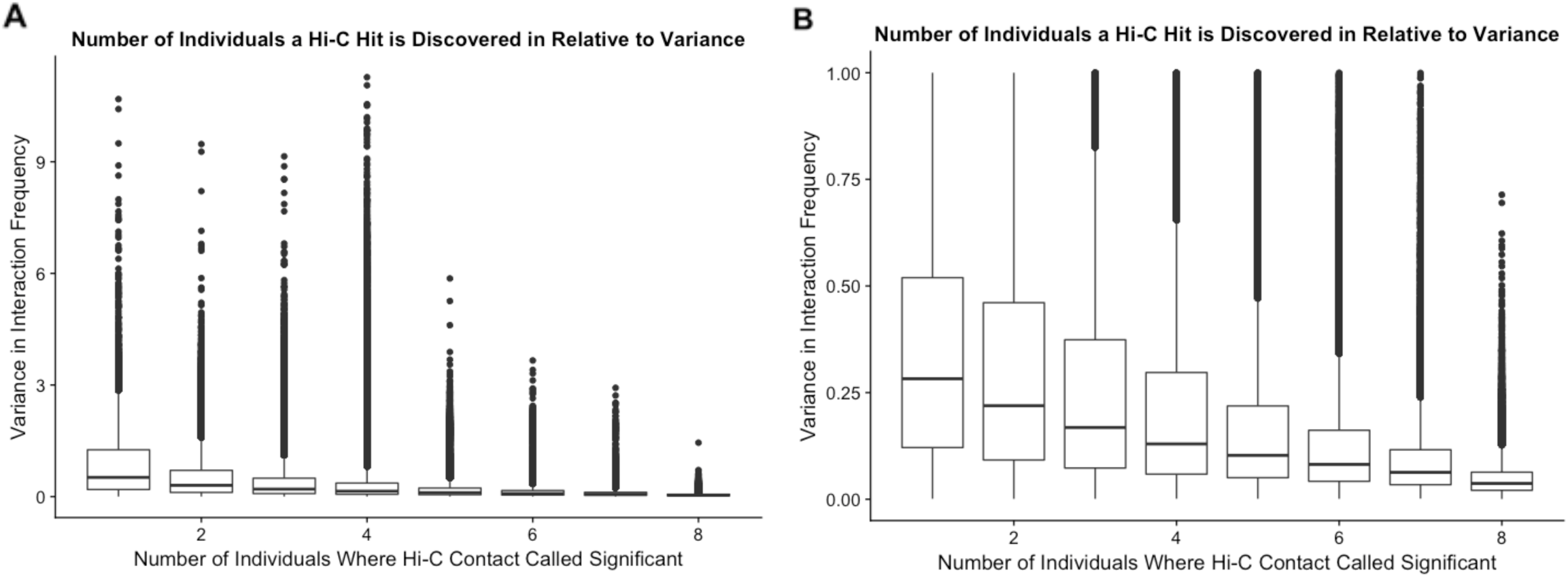
Variance in interaction frequency as a function of the number of individuals in which a significant interaction is independently discovered. **A:** Boxplots of variance in contact frequency across all 8 individuals on the y-axis, binned by the number of individuals in which an interaction is independently called significant on the x-axis. **B:** Same as A, but zoomed in on the y-axis to visualize finer-scale variation.

**Figure S2.**
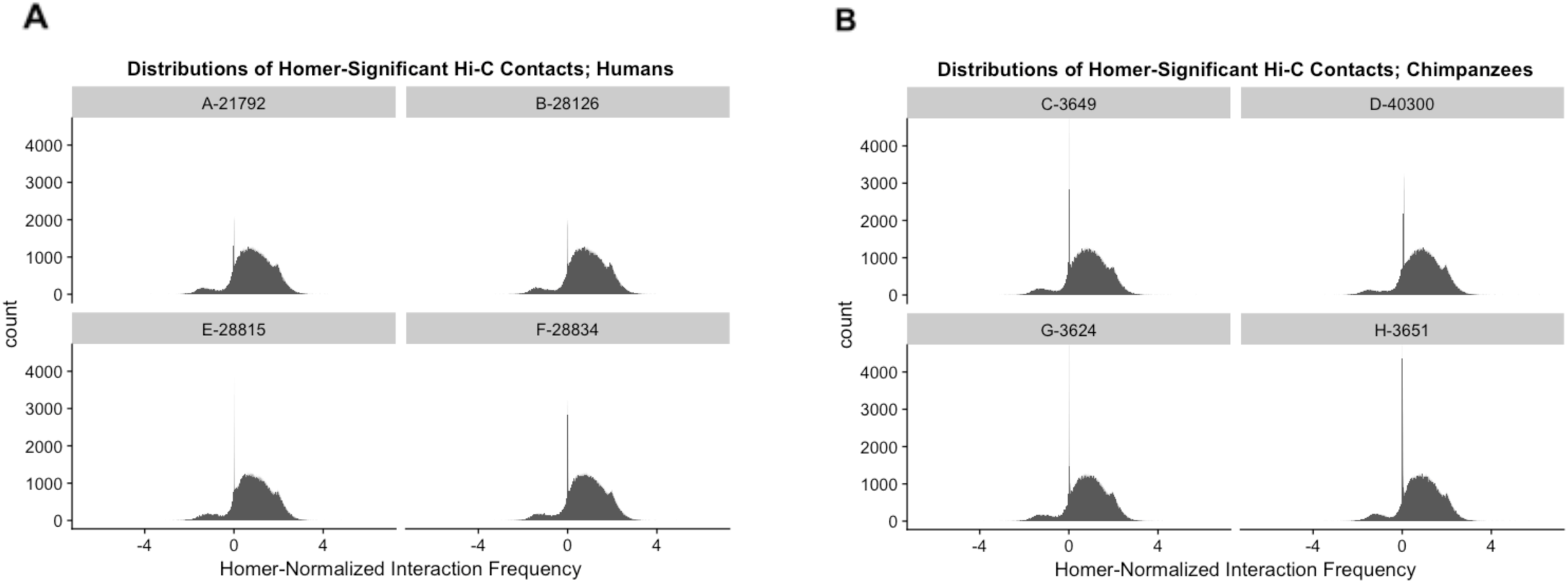
Distributions of HOMER-normalized interaction frequencies are remarkably similar across species. **A:** Histogram of log2(observed/expected) HOMER-normalized interaction frequencies in all four human samples used in this study. **B:** Same as in A, but in chimpanzees.

**Figure S3.**
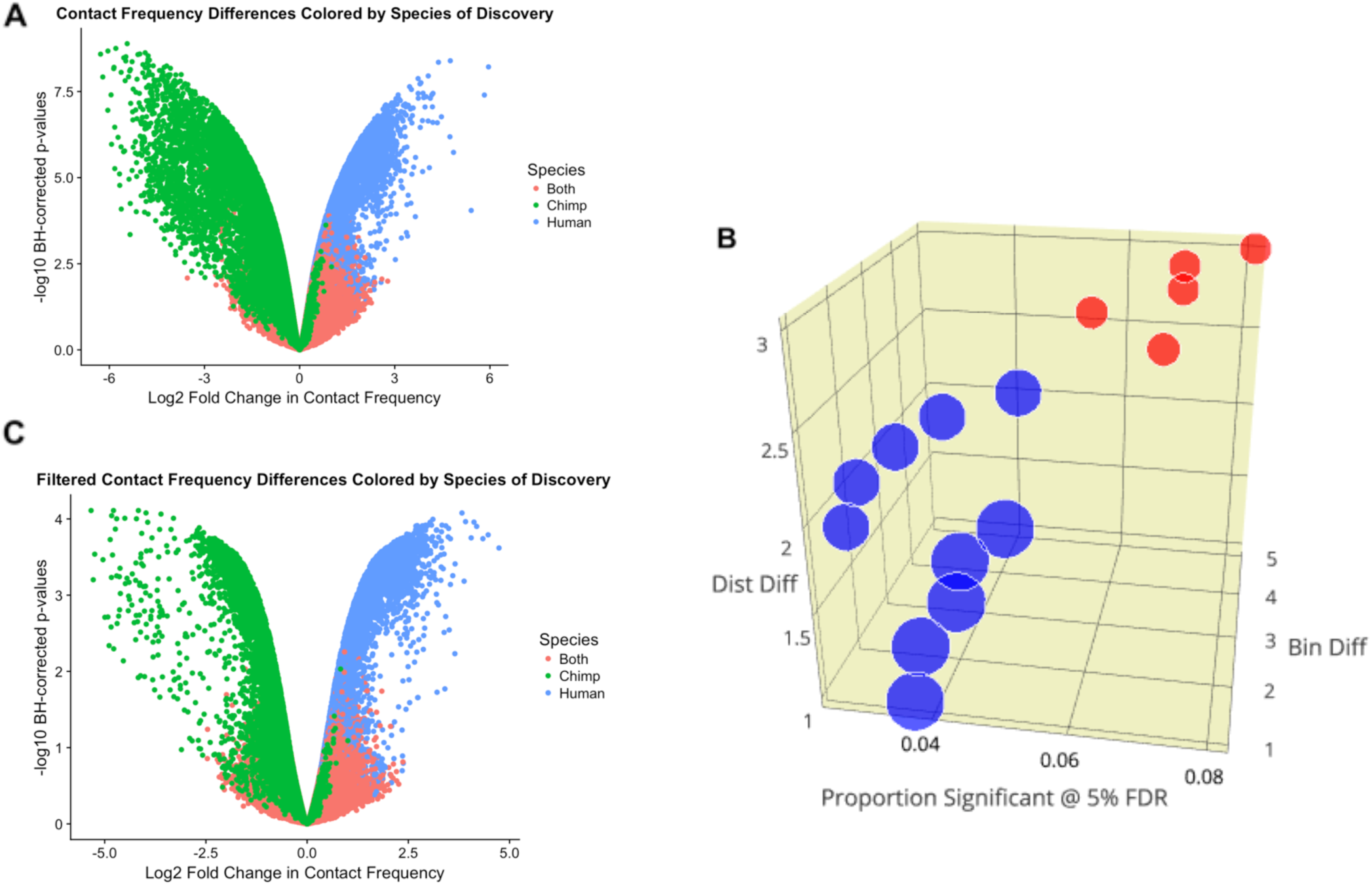
Volcano plot asymmetry quality control. **A:** Volcano plot showing log2 fold change in interaction frequency on the x-axis, and Benjamini-Hochberg corrected –log10 p-values on the y axis. Hits are colored by the species in which they were discovered as significant in HOMER. This plot shows data only filtered for independent discovery in at least 4 individuals. **B:** 3-dimensional plot of sets of Hi-C contacts, with proportion of contacts significant at 5% FDR from linear modeling of interaction frequency on species on the x-axis. Contacts are binned by mate-pair distance differences on the y-axis, and bin size differences on the z-axis. Circle size is proportional to the size of the set of Hi-C contacts falling into each criteria. Red indicates that the data was filtered out after this step, and blue indicates that the data was retained for further analysis. **C:** Volcano plot as in A, but after removing contacts with large mate-pair distance differences across the species induced by liftOver.

**Figure S4.**
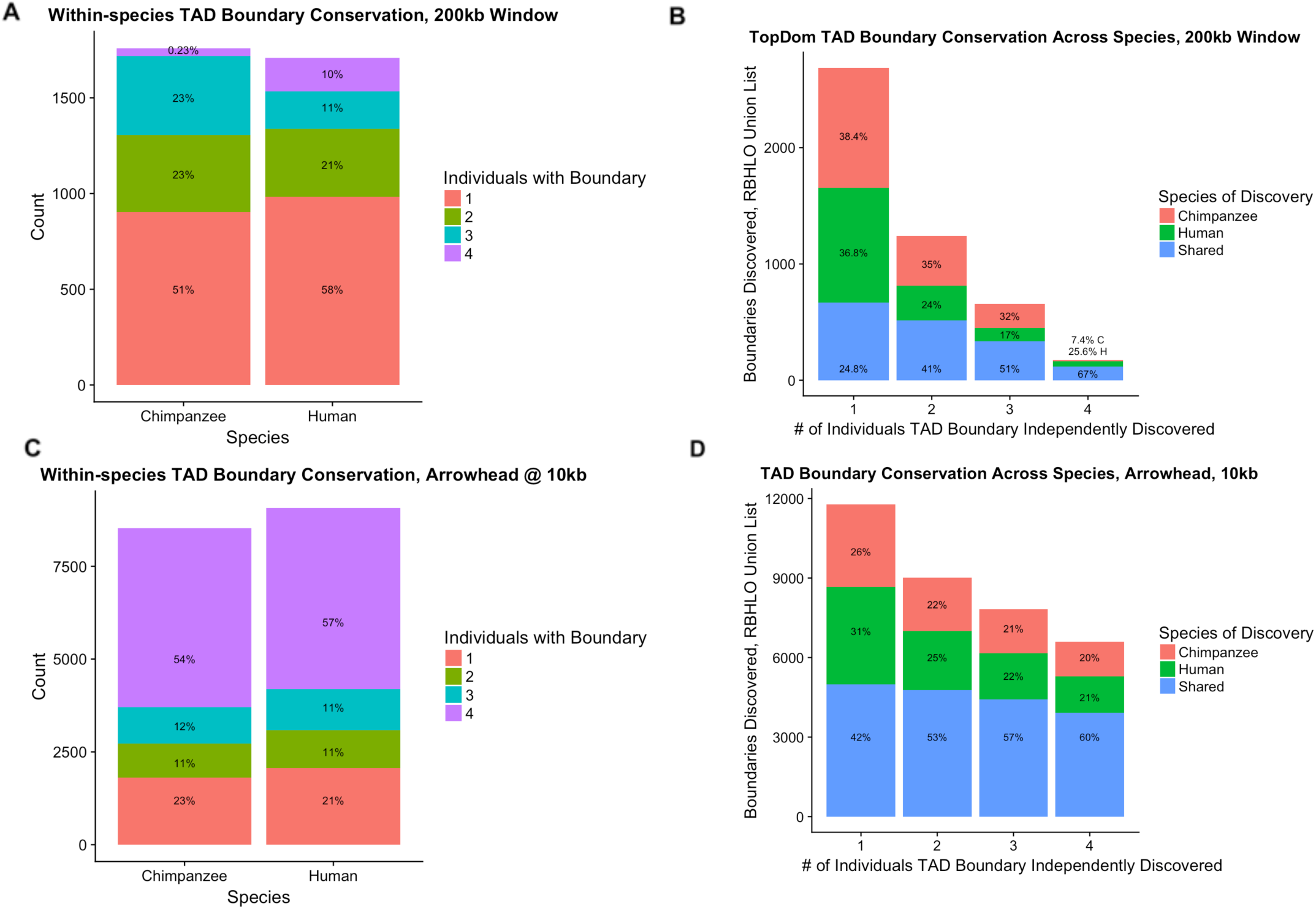
Higher-order chromosomal structure differs substantially between humans and chimpanzees, individual-level data. **A:** Stacked bar chart showing the number of TAD boundaries at a 200kb window that are shared within individuals from each species. In both humans and chimpanzees, approximately half of the TAD boundaries we discover are only found in a single individual. **B**: Similar to A, stacked bar charts with number of individuals (within-species) a hit was discovered in on the x-axis and total boundaries discovered on the y-axis. In this case, however, the union list of boundaries considered is a list across both species, after reciprocally lifting over each species’ individual TAD boundaries to the other genome’s coordinates. When doing no filtering for robustness of boundaries (i.e. when considering all boundaries discovered in at least a single individual), the shared proportion of TAD boundaries is ~25%. Conditioning upon finding a TAD boundary in at least two individuals within a species increases this percentage to 41%. **C:** Same as in A, but this time, TADs were inferred on matrices created in Juicer (KR-normalization) using Arrowhead at 10kb [53]. In both humans and chimpanzees, approximately half of the TAD boundaries we discover are found in all four individuals. **D:** Same as in B, using the Arrowhead analysis described in C. When doing no filtering for robustness of boundaries (i.e. when considering all boundaries discovered in at least a single individual), the shared proportion of TAD boundaries is higher than that seen in the HOMER data (Figure 3B), at 42%. Conditioning upon finding a TAD boundary in at least two individuals within a species increases this percentage to 53%.

**Figure S5.**
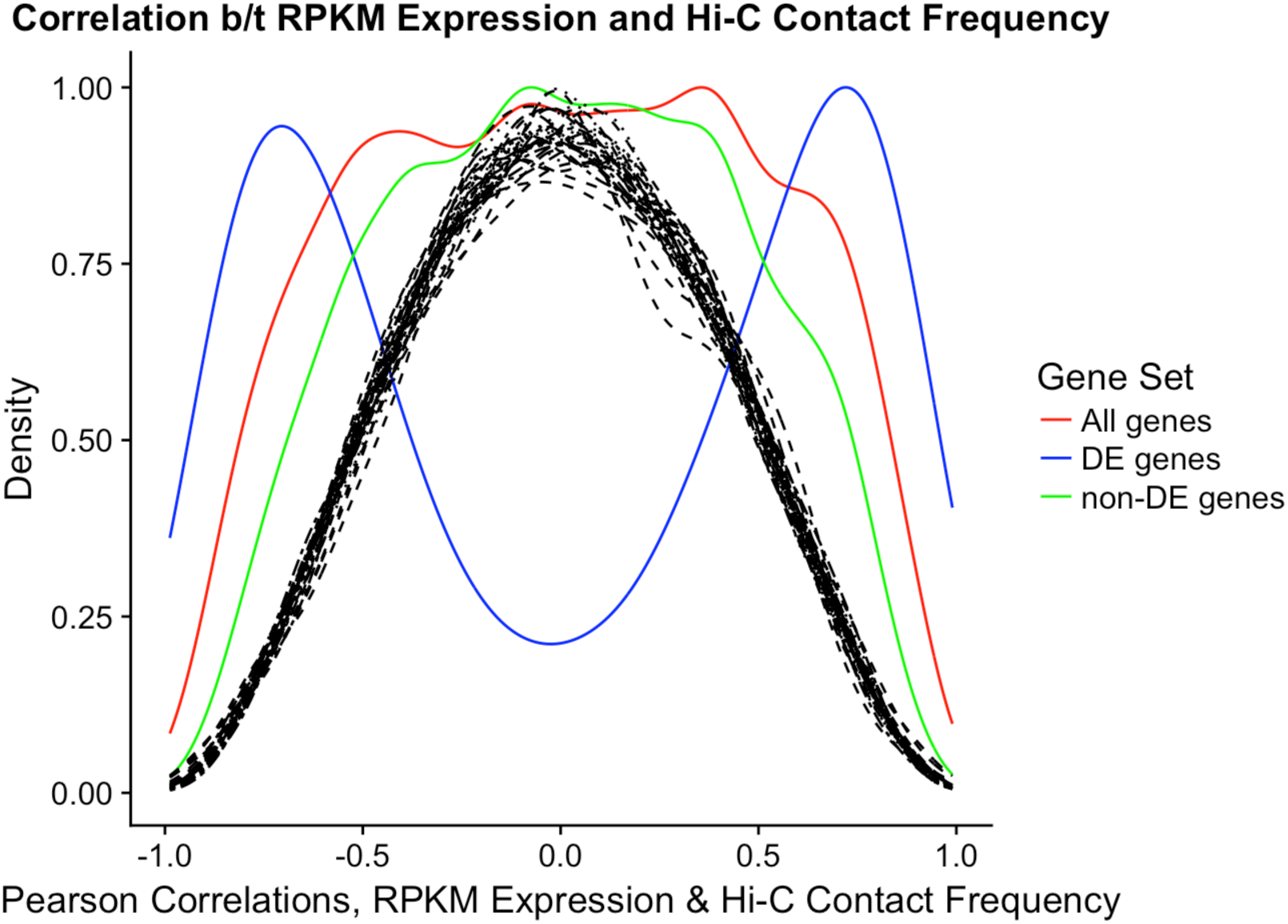
Correlations between Hi-C and expression. Density of Pearson correlations between RPKM expression values and log2 HOMER-normalized contact frequencies. Solid lines indicate different sets of the observed data and dotted lines represent 10 permutations of the data. The Hi-C contact frequency is that with the minimum FDR from linear modeling of contact frequency on species. The strong bimodal distribution of correlations between expression and contact suggests many instances where a contact difference between the species can lead to an increase (enhancer) or decrease (suppressor) of expression in the species where the contact is stronger.

**Figure S6.**
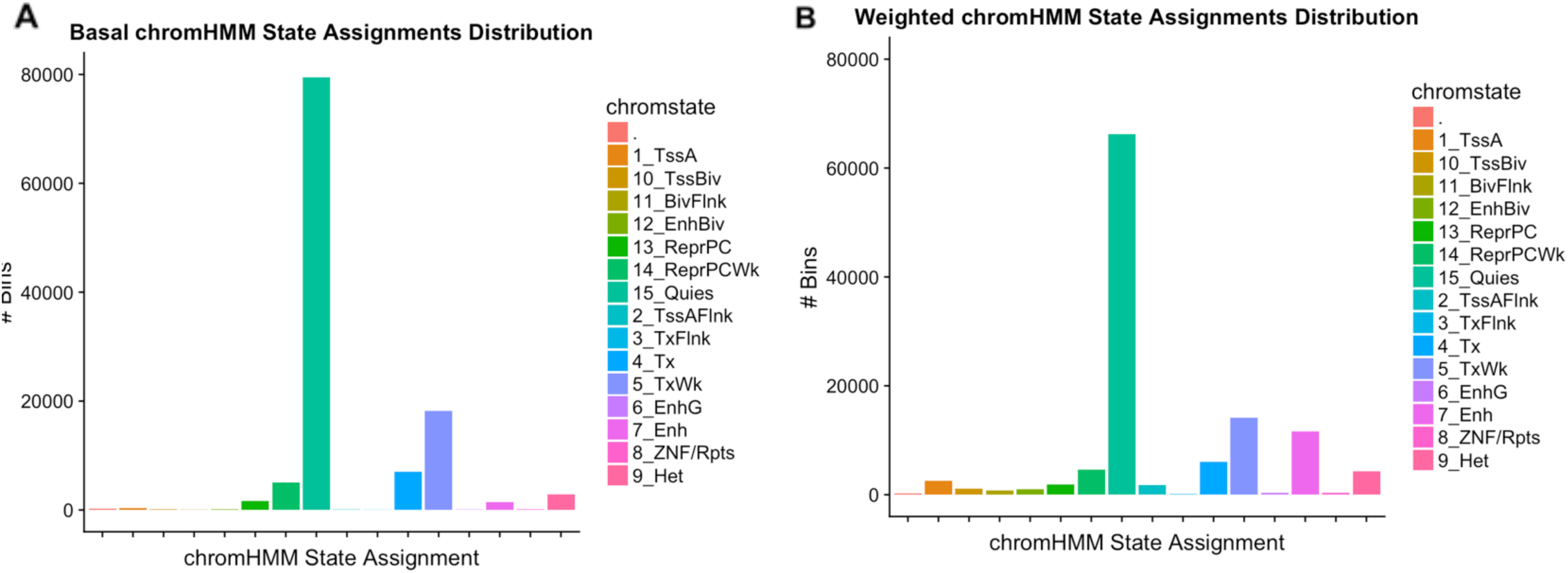
Using a weighting scheme for chromHMM annotations increases the proportion of transcriptional and enhancer-like annotations. **A:** Histogram showing the number of Hi-C loci in each chromHMM annotation (x-axis) using maximum base pair overlap to assign each locus to a state. In the legend, “.” denotes that no annotation was found for a given bin. The number of Hi-C bins in our final list of orthologous contacts that were assigned to a given category is shown on the y-axis. (TssA-Active TSS, TSSBiv-Bivalent/Poised TSS, BivFlnk-Flanking Bivalent TSS/Enh, EnhBiv-Bivalent Enhancer, ReprPC-Repressed PolyComb, ReprPCWk-Weak Repressed PolyComb, Quies-Quiescent/Low, TssAFlnk-Flanking Active TSS, TxFlnk-Transcription at gene 5’ and 3’, Tx-Strong transcription, TxWk-Weak transcription, EnhG-Genic Enhancers, Enh-Enhancers, ZNF/Rpts-ZNF genes and repeats, Het-Heterochromatin) **B:** Same as in A, only here, we assigned annotations after weighting chromHMM elements’ overlaps with Hi-C loci by the reciprocal of their mean overlap in all our loci. This approach increases the number of 10kb Hi-C bins that are assigned to chromHMM annotations associated with transcriptional and enhancer activity (i.e. TssA, TssBiv, TssAFlnk, EnhG, Enh).

**Figure S7.**
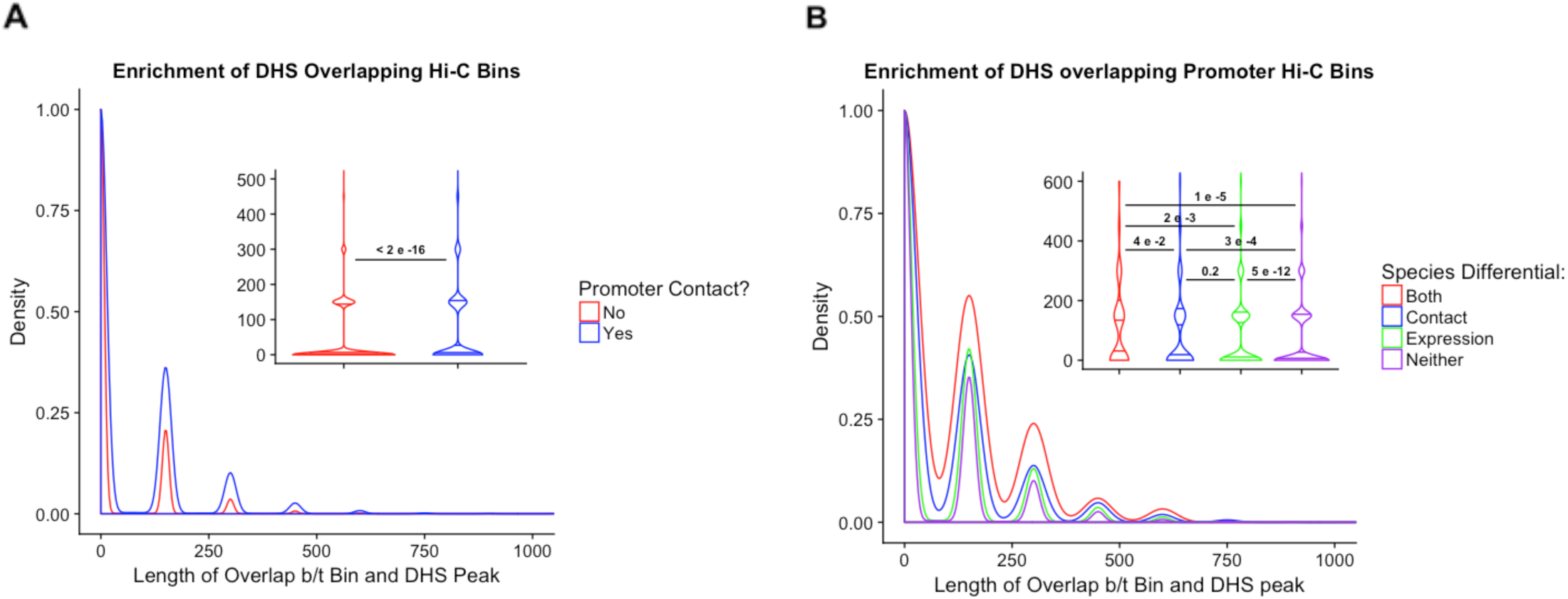
Our significant Hi-C loci are enriched for DHS. **A:** Density distribution of the base pair overlap between DHS peaks downloaded from ENCODE and our Hi-C loci. Plot is split between Hi-C loci that contact a promoter vs. those that do not. Inlay is a violin plot of the same distributions, with lines and numbers indicating pairwise t-tests of the mean, and their corresponding significance levels. **B:** Same as in A, but this time, only considering Hi-C loci involving contact with a promoter, and separating contacts into 4 classes: those that show differential contact between species, those that show differential expression between species, those that show both, and those that show neither.

**Figure S8.**
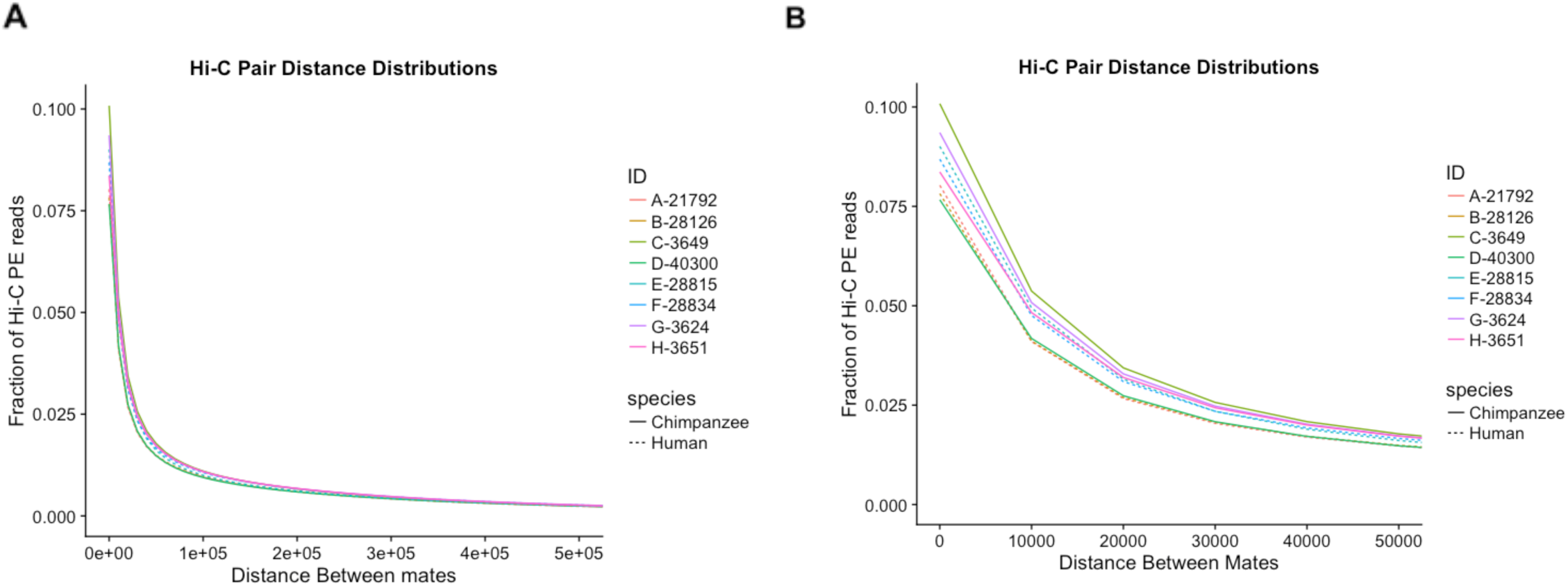
Human Hi-C libraries and chimpanzee Hi-C libraries are comparable. **A:** Fraction of paired-end (PE) reads (y-axis) representing mate pairs at different distances (x-axis), colored differently for each individual library (ID). Dotted lines represent humans, and solid lines represent chimpanzees. **B:** Zoom of A on the x-axis to show the degree of separation between libraries on a smaller scale. Human libraries and chimpanzee libraries do not seem to be clustering separately here.

**Figure S9.**
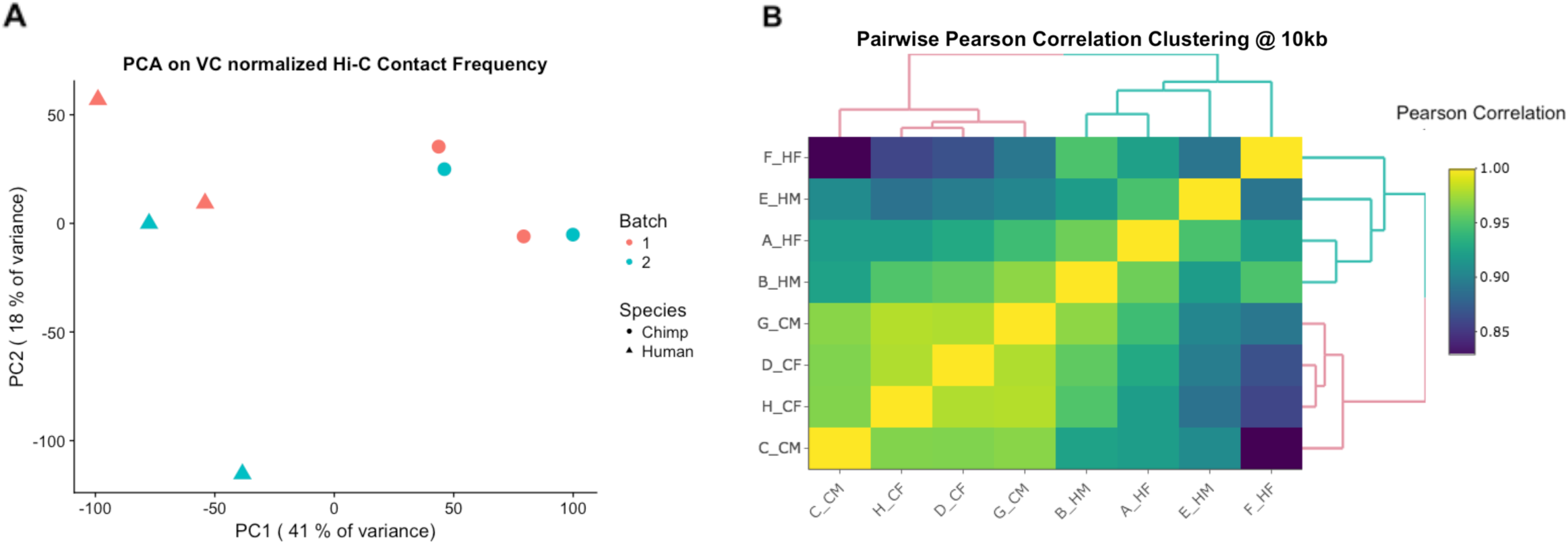
Regulatory landscapes cluster by species, Juicer. **A: Principal components analysis (PCA)** of Juicer vanilla coverage (VC)-normalized interaction frequencies for the union of all significant hits between humans (triangles) and chimpanzees (circles). PC1 captures ~41% of the variance, and is correlated with species (r = 0.68 and P = 0.01 in regression of PC1 by species). **B: Unsupervised hierarchical clustering** at 10 kb of the pairwise correlations (Pearson’s r^2^) between Juicer VC-normalized interaction frequencies. The first letter in the labels demarcates the species (H for human and C for chimpanzee), and the following symbols indicate sex (male, M or female, F) and batch (1 or 2).

**Figure S10.**
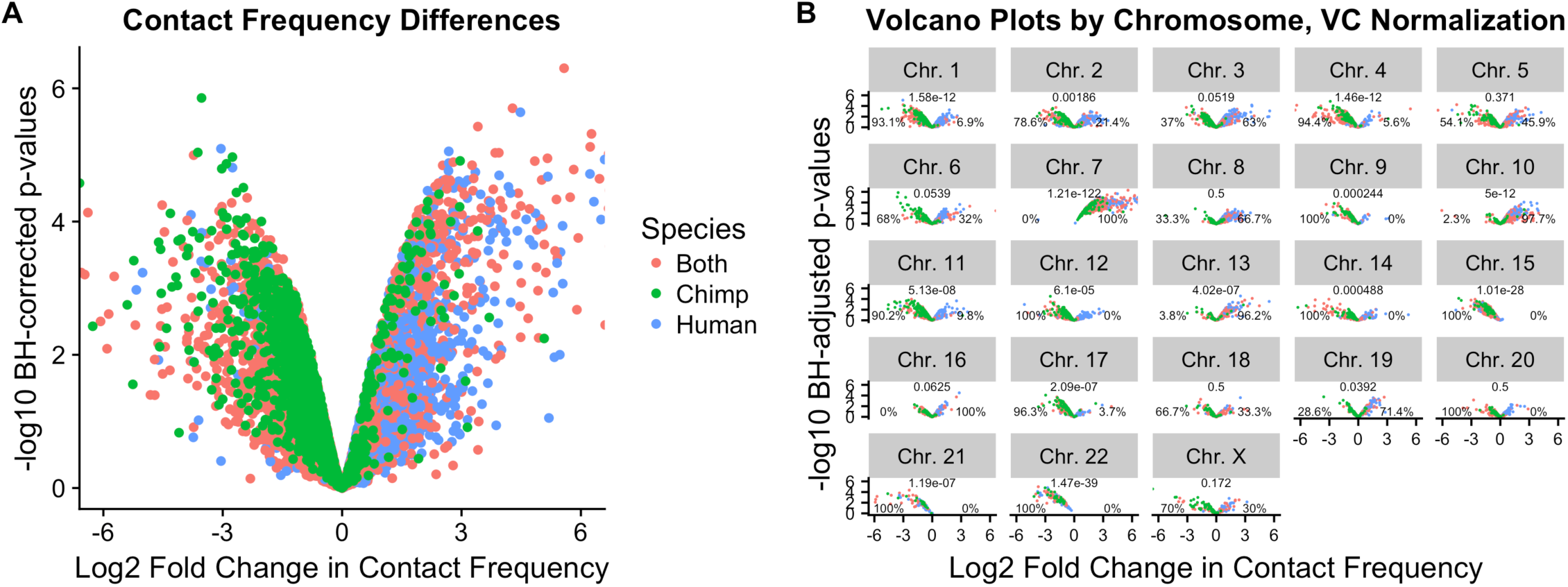
Linear modeling reveals large-scale chromosomal differences in contact frequency, Juicer. **A Volcano plot** of log2 fold change in contact frequency between humans and chimpanzees (x-axis) against Benjamini-Hochberg corrected –log10 p-values (y-axis),after filtering out data affected by putative errors in orthology assignment. Because humans were coded as a 1 in the linear model, negative values represent pairs with decreased contact frequency in humans compared to chimpanzees, and positive values represent pairs with increased contact frequency in humans. **B: Per-chromosome volcano plot** using the same legend as in A. Numbers above each panel represent p-values from a binomial test under the expectation of significant hits being evenly distributed across the species. Percentages to the left and right of each plot represent the percentage of significant hits that are stronger in chimpanzees and humans, respectively. Of note is that many of the same chromosomal asymmetries in contact strength observed here are in the same chromosomes as those observed in the Homer-normalized data (Figure 2).

**Figure S11.**
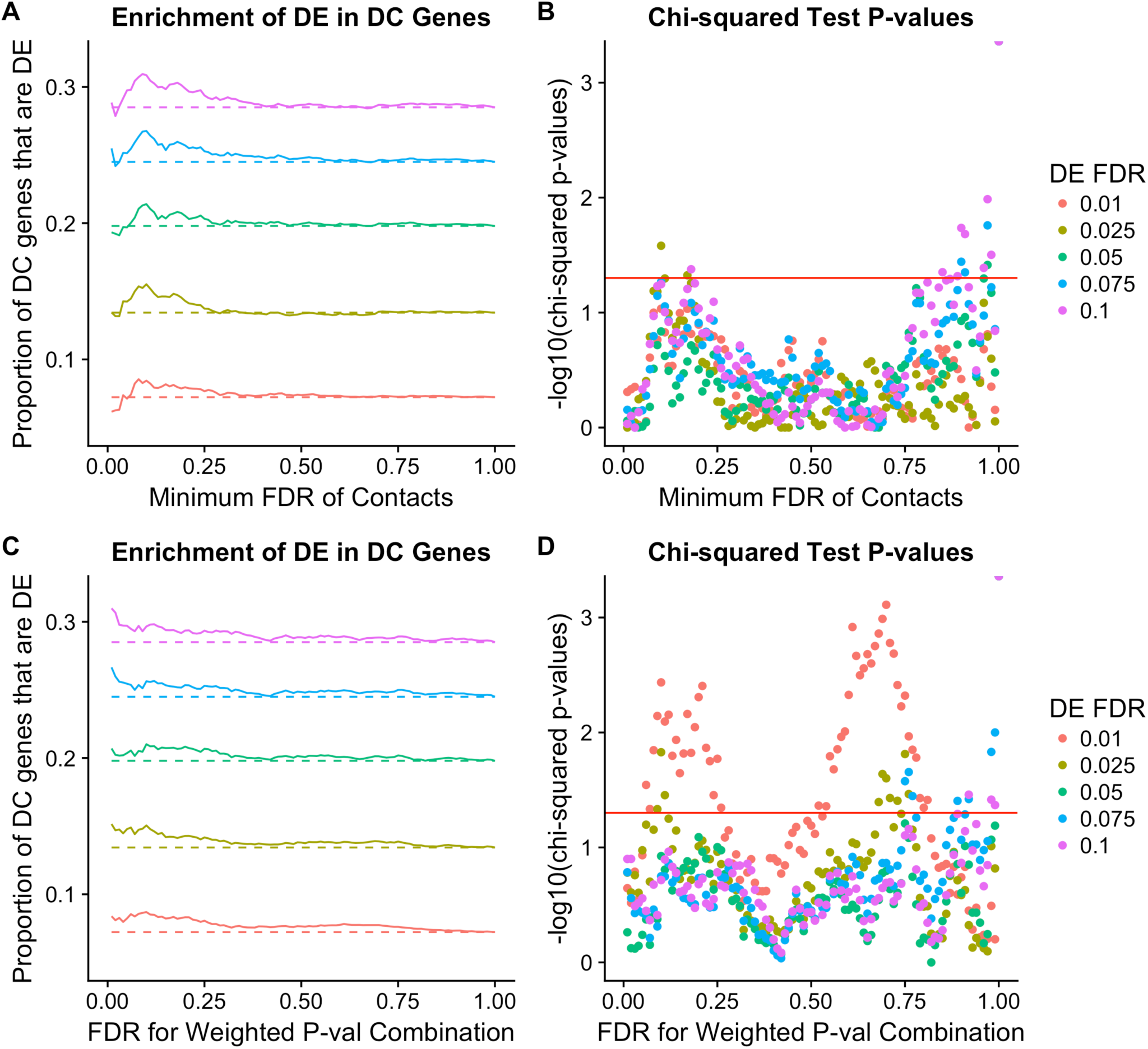
Differentially expressed genes show enrichment for differential Hi-C contacts, Juicer. **A:** Enrichment of differential expression (DE) in genes with differential Hi-C contact frequencies (DC). The proportion of DC genes that are significantly DE (y-axis) is shown across a range of DC FDRs (x-axis). Colors indicate different DE FDR thresholds and dashed lines indicate the expected proportion of DC genes that would be DE based on conditional probability alone. **B:** Chi-squared significance tests for the enrichments in A. –log10 chi-squared test p-values for enrichment of DC in DE genes (y-axis) is shown for a range of DC FDRs (x-axis). If a gene had multiple DCs, we plotted only the contact that had the minimum FDR. As in A, colors indicate different DE thresholds. A horizontal red line is drawn where p=0.05. The strongest signal of enrichment is observed at 2.5% DE FDR and 20% DC FDR, with *P* = 0.01. This is comparable to enrichments found in the HOMER-normalized set of data, where the strongest signal was seen at 2.5% DE FDR and 10% DC FDR (Figure 4A). **C:** Same as in A, but this time, a weighted p-value combination technique [62] was used to integrate each Hi-C bin’s DC FDR across all of its contacts. **D:** Same as in B, but for the weighted p-value combination instead of the minimum FDR contact.

**Figure S12:**
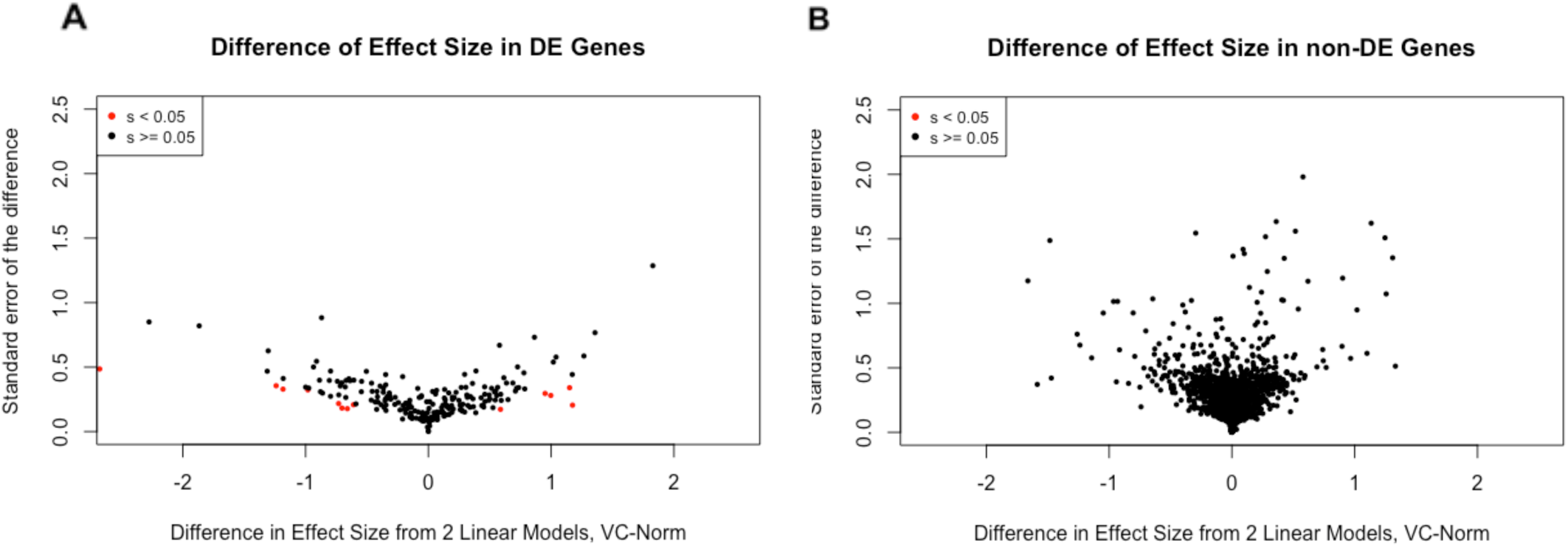
Gene expression variance is explained by chromatin contacts for 6% of DE genes, Juicer. **A:** Volcano plot of the differences in DE effect size between models before and after conditioning on contact frequency (x-axis) plotted against the standard error of the difference (y-axis). Adaptive shrinkage was used to evaluate significance of the differences in effect size. Amongst DE genes, 6% (13/236) of genes showed a statistically significant and sign-confident effect of Hi-C contacts on expression levels (s-value ≤ 0.05). **B:** Same as in A, but performed on the set of 1,394 non-DE genes. No genes showed a statistically significant difference in effect size between the two models.

**Figure S13.**
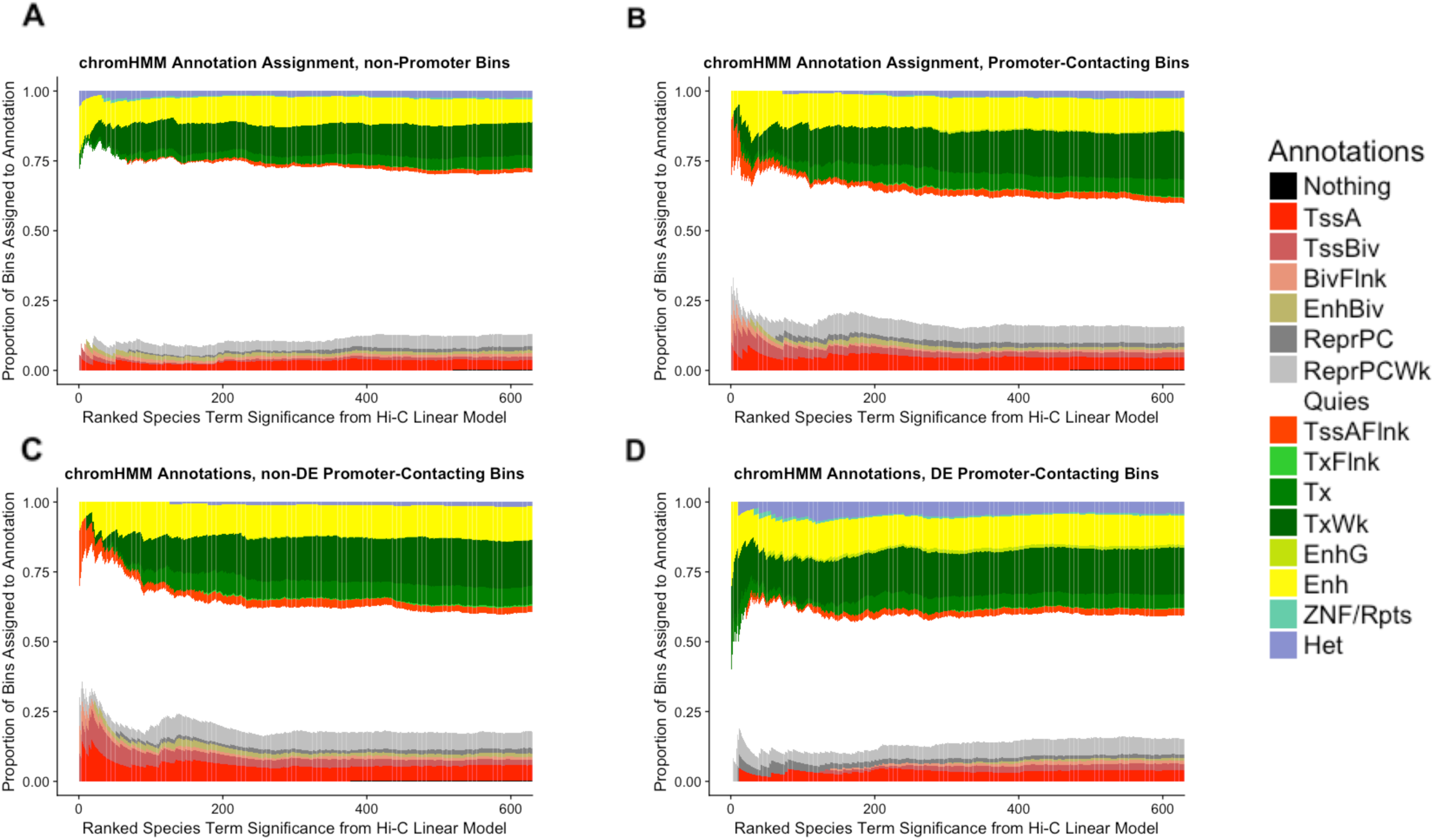
Dynamics of chromHMM state among significant Hi-C contacts, Juicer. In A-D, Hi-C contact loci are ranked in order of decreasing significance (x-axis), with species-specific contacts (i.e. lower FDR for our species term from linear modeling) on the left, decreasing in significance towards the right. The y-axis shows cumulative proportion of chromHMM annotation assignments for all Hi-C loci at the given rank or lower. (TssA-Active TSS, TSSBiv-Bivalent/Poised TSS, BivFlnk-Flanking Bivalent TSS/Enh, EnhBiv-Bivalent Enhancer, ReprPC-Repressed PolyComb, ReprPCWk-Weak Repressed PolyComb, Quies-Quiescent/Low, TssAFlnk-Flanking Active TSS, TxFlnk-Transcription at gene 5’ and 3’, Tx-Strong transcription, TxWk-Weak transcription, EnhG-Genic Enhancers, Enh-Enhancers, ZNF/Rpts-ZNF genes and repeats, Het-Heterochromatin) **A:** chromHMM annotation assignment for non-promoter contacting bins. **B:** chromHMM annotation assignment for all promoter-contacting bins. **C:** chromHMM annotation assignment for non-DE promoter-contacting bins. **D:** chromHMM annotation assignment for DE promoter-contacting bins.

**Figure S14.**
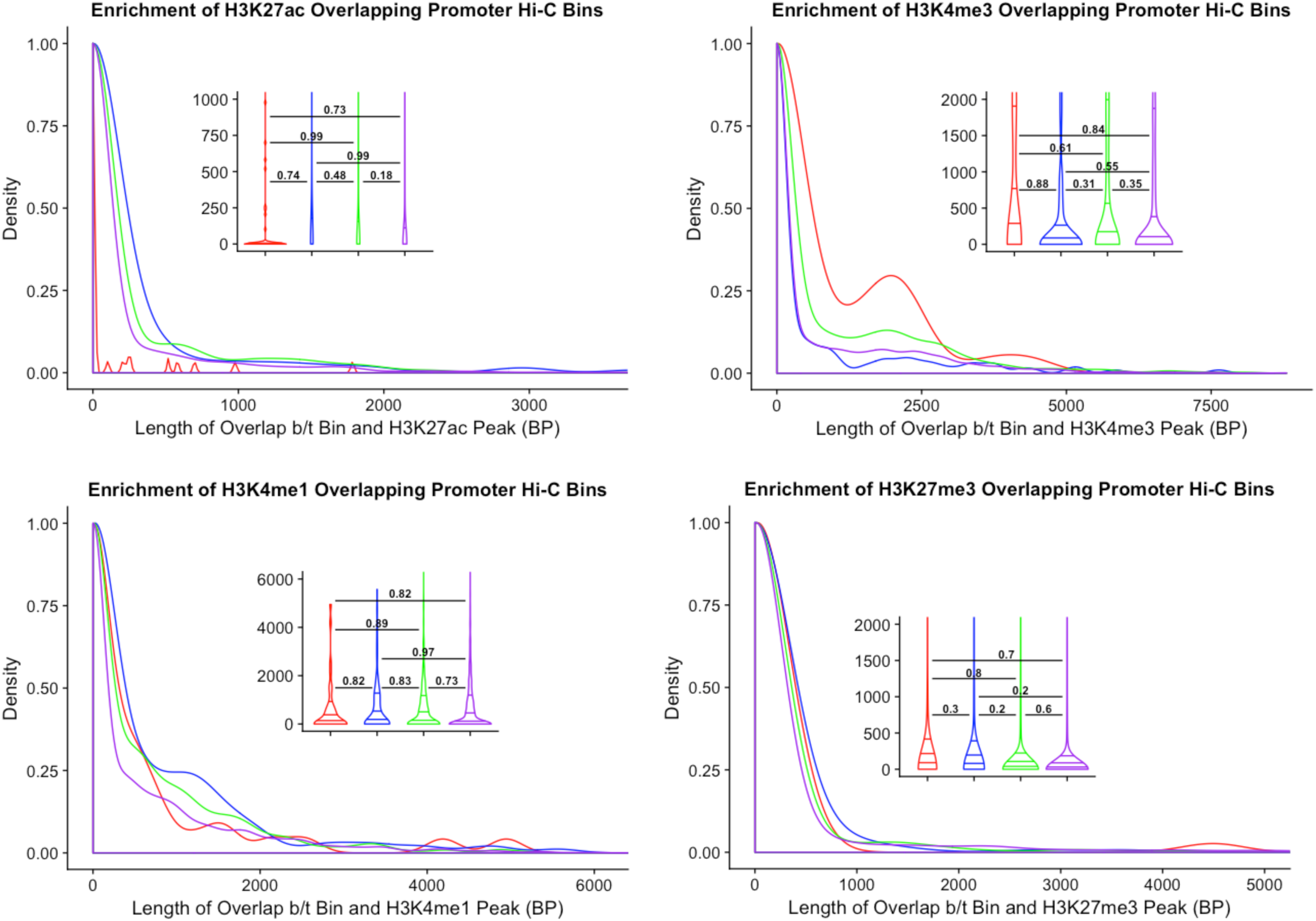
Overlap of activating and repressive histone marks among Hi-C contacts, Juicer. A-D show density plots of the base pair overlap between different classes of Hi-C contact loci and various histone marks. All histone mark data were obtained from ENCODE in experiments carried out in human iPSCs (A-C) or hESCs (D). We grouped contacts into 4 classes, indicated by color: those that show differential contact between species, those that show differential expression between species, those that show both, and those that show neither. We used pairwise t-tests to compare differences in the mean overlap among the four classes of Hi-C loci. Inlaid violin plots show the significance level of each t-test, with horizontal lines connecting the two groups used for comparison. Lines inside each violin represent the 25^th^, 50^th^, and 75^th^ percentiles of the data. Unlike in the HOMER-normalized data, we do not observe statistically significant differences in overlaps with H3K27ac between different locus classes. This may reflect the previous observation that the hiccups algorithm for assigning statistical significance of loops in Hi-C data is much more conservative than HOMER’s significance calling method [93]. Additionally, in our analyses, the number of promoter-contacting bins we observe as statistically significant is an order of magnitude smaller in the Juicer data (~1k) than in the HOMER data (~11k). The significantly smaller set available here may explain why we do not see the same epigenetic mark enrichment dynamics as we observed in the HOMER data (Figure 7). **A:** Density plot of base pair overlap between Hi-C contact loci and the active histone mark H3K27ac. **B:** Same as in A, but for the active histone mark H3K4me3. **C:** Same as in A and B, but for the active histone mark H3K4me1. **D:** Same as in A-C, but for the repressive histone mark H3K27me3.

**Figure S15:**
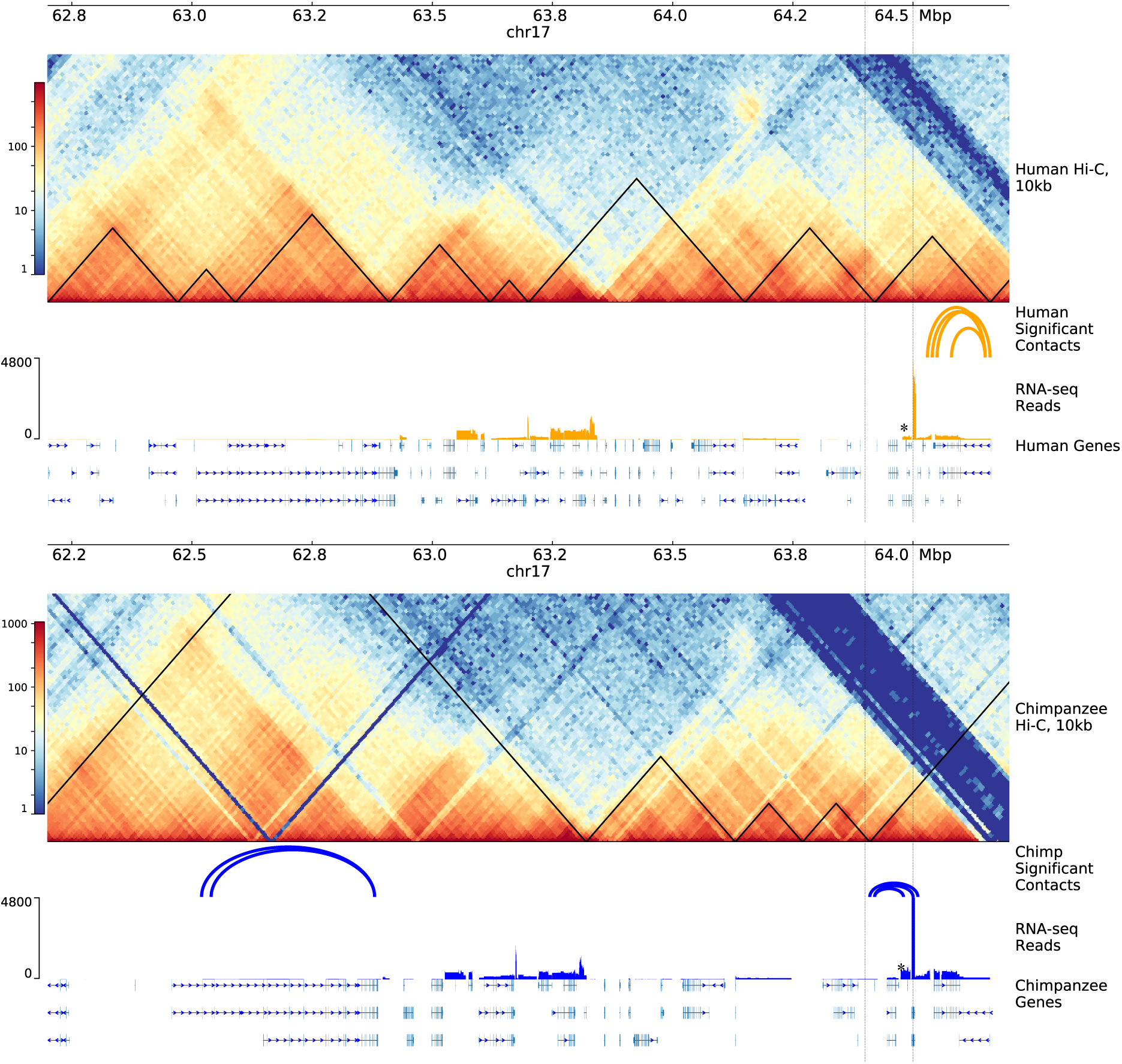
Broad view of gene expression and Hi-C contact. PyGenomeTracks plot [70] of a 2mb syntenic region between the species, zoomed out from the region displayed in Figure 6 (dotted black lines). Humans are displayed on the top panel and chimpanzees on the bottom panel, with Hi-C matrices at 10kb resolution and TADs overlaid as black lines. The first track underneath Hi-C matrices in both panels is an arc diagram displaying the differential contacts observed to be stronger in the given species in our dataset. Shown beneath this are raw RNA-seq reads that mapped to orthologous exons between the species, followed by a track of gene predictions (*POLG2* marked by an asterisk).

